# Nematode-Trapping Devices of *Arthrobotrys oligospora* is an Iron Storage Phenotype Adapted to Temperature Changes

**DOI:** 10.1101/2024.07.25.605209

**Authors:** Jiao Zhou, Qunfu Wu, Li Wu, Ling Li, Songhan Xue, Junxian Yan, Zumao Hu, Xuemei Niu

## Abstract

Under low-nutrient conditions, *Arthrobotrys oligospora* and other NTFs can differentiate their mycelia into specialized trapping devices for capturing prey as their nutritional source. Using energy-dispersive X-ray spectroscopy (EDX) in conjunction with transmission electron microscopy (TEM), we identified that the characteristic electron-dense bodies in trapping devices contained more iron than vacuoles and mitochondria. Meanwhile, fungal mycelial cells used effective desferriferrichromes for iron chelation and storage. Complex bioassays showed that electron-dense bodies represent a novel type of microbial iron storage particle and trapping devices in *A. oligospora* function as an unprecedented phenotypic system for iron storage. Unexpectedly, all NTFs lack a crucial Ccc1-mediated vacuolar iron detoxification mechanism, which is conserved in most fungi. Inserting the *Ccc1* gene cloned from yeast into *A. oligospora* significantly reduced formation of trapping devices and inhibited nematicidal activity. Notably, Bayesian relaxed molecular clock analysis indicated that the loss of Ccc1-mediated vacuolar iron storage occurred during the Late Paleozoic Ice Age, while the origin of the trapping devices and the acquisition of desferriferrichrome biosynthesis were strongly associated with significantly elevated temperatures. Temperature bioassays demonstrated that the formation of trapping devices is highly temperature-dependent, with free iron content in mycelial cells being inversely proportional to temperature, consistent with that *A. oligospora* is sensitive to high temperatures and fails to grow above 30°C. Our findings revealed that global temperature fluctuations are a crucial driver of the genetic evolution of NTFs, as a catalyst for the origin of trapping devices, which are a novel phenotypic indicator of eukaryotic iron overload.

**Author summary:** We found that a unique group of carnivorous fungi has evolved specialized trapping devices to sequester excess iron, compensating for the absence of the crucial Ccc1-mediated vacuolar iron detoxification pathway, which is conserved in most fungi. Furthermore, we report for the first time that elevated temperature is a non-trophic factor that induces iron overload in eukaryotes, as iron content and composition in fungal mycelia are negatively correlated with temperature. Our findings suggest that nematode-trapping fungi could serve as a potential eukaryotic model for investigating the dynamic regulation mechanisms of iron homeostasis, which could contribute to the development of therapies for iron overload-related diseases in humans. In humans, iron overload leads to tissue damage, particularly in the cardiovascular system. It has long been assumed that iron overload occurs when iron intake is increased over an extended period, either through repeated red blood cell transfusions or enhanced absorption from the gastrointestinal tract. Caucasians are particularly susceptible to iron overload and the complications of hemochromatosis due to a higher incidence of mutations in the homeostatic iron regulation gene within this population. The discovery of eukaryotes exhibiting an iron overload phenotype when exposed to heat holds significant implications for developing treatments and strategies for human iron overload disorders.

## Introduction

Nematode-trapping fungi (NTFs), which belong to the phylum Ascomycota and form a monophyletic group within the class Orbiliomycetes, are capable of differentiating their mycelia into specialized trapping devices under nutritionally poor conditions, including adhesive networks, adhesive knobs, and constricting rings, to capture nematodes, small animals, and protozoans as their primary nutritional sources [1–3]. The formation of these trapping devices serves as a diagnostic indicator of the transition of NTFs from a saprophytic to a predatory lifestyle [4]. The flexible lifestyle and adaptability of NTFs make them a promising candidate for controlling parasitic nematodes in plants and animals [5]. *A. oligospora* is one of the most extensively studied NTFs, found in diverse regions across the globe, from tropical to Antarctic zones and from terrestrial to aquatic ecosystems. In the presence of nematodes, *A. oligospora* under poor nutritional conditions can form complex, three-dimensional adhesive networks to capture them. Previous studies have shown that *A. oligospora* also detect and respond to ammonia, certain amino acids, and other nematode-related molecules, which are small, evolutionarily conserved signaling compounds secreted by many species of soil-dwelling nematodes and bacteria. These molecules act as signals to recognize prey and trigger the formation of trapping devices [6, 7]. *A. oligospora* has long served as an excellent model system for understanding the evolution of fungi and their interactions with nematodes [5]. However, few studies have addressed the role of specific environmental factors in the induction of trapping devices, with most research emphasizing that nutrient deficiency is essential for trapping device formation.

The ability to sequester excess elements for storage is essential for survival and fitness in all organisms. In eukaryotes, a key mechanism for iron storage involves ferritin or siderophore-mediated sequestration, which forms ferric-containing complexes [8–10]. In pathogenic fungi, siderophores are often considered virulence factors, making siderophore biosynthesis a potential target for antifungal chemotherapy against *Candida albicans*, as the underlying biochemical pathways are absent in human cells [11]. Desferriferrichrome, the first known fungal siderophore and ferric chelating agent, was isolated from the corn smut fungus *Ustilago sphaerogena* in 1952. It is widely distributed in fungi as a well-known siderophore for iron uptake and storage [12, 13]. *A. oligospora* also produces desferriferrichrome through two genes, *sidA* and *NRPS*, which encode L-ornithine N(5)-oxygenase and a nonribosomal peptide synthetase, respectively [14]. Recent studies have shown that deficiencies in *sidA* or NRPS not only result in the loss of desferriferrichrome in *A. oligospora*, but also trigger the mass production of trapping devices in the Δ*sidA* and Δ*NRPS* mutants, compared to WT [14]. Furthermore, the Δ*sidA* mutant, which lacks all desferriferrichrome precursors, develops significantly more trapping devices than the Δ*NRPS* mutant, which retains some desferriferrichrome precursors capable of sequestering iron [14]. Surprisingly, most of the iron-induced trapping devices failed to capture nematodes. Supplementing with desferriferrichrome partially restored the nematode-capturing ability of the trapping devices, but significantly inhibited their formation. These unexpected findings reveal, for the first time, that iron is an abiotic factor that induces trapping device formation, and that the formation of trapping devices is not related to the nematicidal activity of *A. oligospora* [14].

Notably, in the fungal kingdom, vacuolar iron storage—rather than siderophore-mediated storage—is the primary mechanism for iron detoxification, owing to its capacity to store excess iron [12, 15]. The vacuolar iron transporter Ccc1 is highly conserved across fungi and is essential for importing iron into the vacuole to prevent toxic iron overload in the cytosol [16–18]. Ccc1-mediated vacuolar iron detoxification is particularly crucial for pathogenic fungi, which must adapt to grow inside warm-blooded hosts [19, 20]. Inactivation of the *Ccc1* gene in the human pathogenic fungus *Aspergillus fumigatus* significantly reduces resistance to iron toxicity [17]. A previous study have shown that the absence of desferriferrichrome leads to significantly elevated ferric ion levels in the Δ*sidA* mutant, indicating that vacuoles in *A. oligospora* fail to store the excess iron resulting from the loss of desferriferrichrome-mediated iron storage [14]. Therefore, the molecular basis for the formation of trapping devices induced by iron overload in the mutant Δ*sidA* warrants further investigation. Moreover, based on molecular clock and fossil evidence, NTFs are estimated to have originated as a result of mass extinctions during the Permian and Triassic periods [1]. However, solid evidence is lacking regarding which specific genes were lost or acquired during these periods and contributed to the development of trapping devices in NTFs.

Previous studies reported that mature trap cells are typically filled with "special" electron-dense bodies, which form during the early stages of trap development and are considered peroxisomal in nature [21, 22]. These electron-dense bodies are absent in normal vegetative cells, which contain "normal" microbodies. However, the chemical properties of these bodies in trap and vegetative cells remain unclear. To investigate this, we examined the role of iron in the contents of electron-dense bodies in the trap cells of *A. oligospora* using transmission electron microscopy (TEM). We then employed energy-dispersive X-ray spectroscopy (EDX) in conjunction with TEM and found that the electron-dense bodies in trap cells contained significantly more iron than vacuoles and mitochondria. Importantly, we observed that desferriferrichrome-mediated iron storage is primarily occurs in vegetative cells and more efficient than electron-dense bodies. The inefficiency of vacuoles for iron storage led us to discover that NTFs lack eight key vacuole-related orthologous genes, including *Ccc1*, *thi4*, *ivy1*, *gnt1*, *vtc5*, *sna3*, *opy2*, and *phm6*. Insertion of a *Ccc1* gene cloned from yeast into *A. oligospora* resulted in decreased trapping devices and nematicidal activity of the fungus. A Bayesian relaxed molecular clock analysis, utilizing 275,133 four-fold degenerate sites from 1,136 single-copy orthologous genes and three fossil calibration points, indicated that the loss of these vacuole-related genes and the origin of trapping devices are linked to the transition from the Late Paleozoic Ice Age to the Permian and Triassic extinctions.

Importantly, we found that the acquisition of desferriferrichrome-mediated iron storage was associated with high temperatures during the Cambrian period, while the loss of Ccc1-mediated vacuolar iron storage was linked to the longest ice age of the Late Paleozoic, rather than to iron levels throughout the Phanerozoic eon. Notably, the origin of trapping devices was also associated with lethally high temperatures during the early Triassic Greenhouse, just after the longest ice age. Phenotypic analysis revealed that *A. oligospora* exhibited optimal growth at 24°C but failed to grow above 30°C. Importantly, elevated temperatures strongly promoted the formation of trapping devices, even in the absence of nematode inducers or iron supplementation. Collectively, our work demonstrates that high-temperature-sensitive nematode-trapping fungi (NTFs), which are deficient in vacuole-mediated iron storage, have evolved trapping devices as a mechanism to store excess iron induced by elevated temperatures.

## Results

### 1. Lack of desferriferrichromes promoted the contents of electron dense bodies in trap cells

We first evaluated the role of desferriferrichromes in trapping device formation using the model nematode *Caenorhabditis elegans* WT within 24 hours, with strains lacking nematodes serving as controls. Remarkably, in the absence of nematode treatment, WT did not develop trapping devices within 24 hours, while the mutant Δ*sidA* spontaneously began to form immature trapping devices (Fig. 1), consistent with previous findings that the absence of desferriferrichromes induces spontaneous trapping device formation [14]. With nematode treatment, WT initiated the formation of immature trapping devices within 4 hours, and the trapping devices matured within 11 hours. In contrast, the mutant Δ*sidA* formed mature trapping devices within 8 hours (Fig. 1). Importantly, nematodes in the Δ*sidA* group were physically destroyed and dissolved within 10 hours, while those in the WT group were not until 16 hours (Fig. 1).

**Fig 1.**
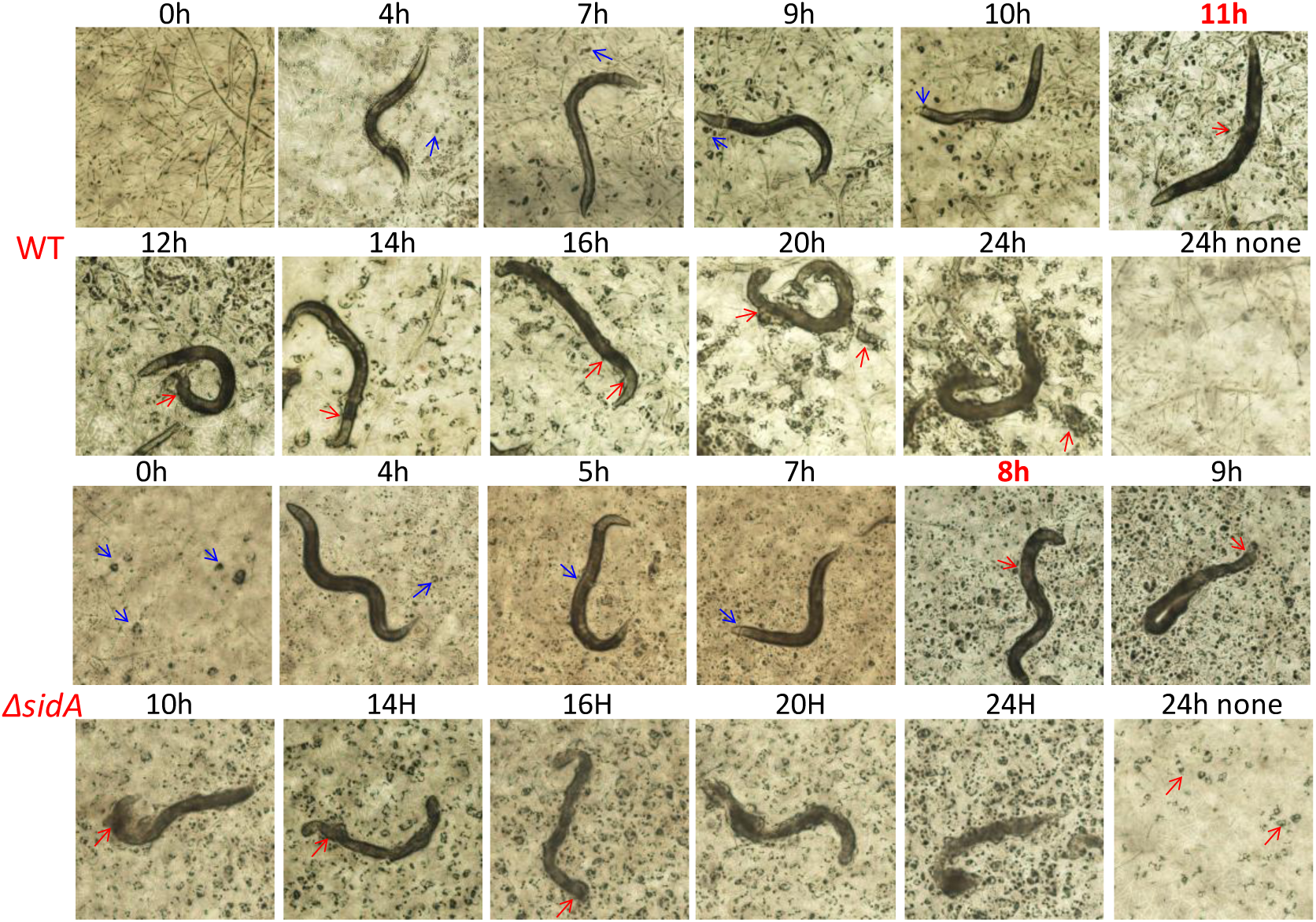
Evaluation of the role of iron-chelating desferriferrichromes in the formation of trapping devices of *A. oligospora*. Comparison of the formation of trapping devices in WT and the mutant *ΔsidA* treated with nematodes within 24 h revealed that the lack of iron-chelating desferriferrichromes promoted the formation of trapping devices and nematicidal activity. Nematodes were physically dissolving at 10h in mutant *ΔsidA* group, but at 16h in WT group. 24h-none: the strain without nematode treatment for 24h. Blue arrow: immature trapping devices. Red arrow: mature trapping devices.

The accumulation of electron-dense bodies is a key diagnostic feature of trap cells, we evaluated the formation of electron-dense bodies in *A. oligospora* WT and the mutant Δ*sidA* treated with nematodes. Transmission electron microscopy (TEM) analysis revealed that the formation of electron-dense bodies occurs in two distinct stages: the first stage involves the formation of numerous small amorphous particles (referred to as small electron-dense bodies), which then aggregate to form larger, tangible electron-dense bodies (Fig. 2a). Notably, the mutant Δ*sidA* accumulates a large number of small electron-dense bodies within 4 hours, while WT requires at least 7 hours (Fig. 2a and S1), suggesting that the absence of the iron-chelating desferriferrichromes accelerates the early stage of small electron-dense body formation in the mutant Δ*sidA*.

**Fig 2.**
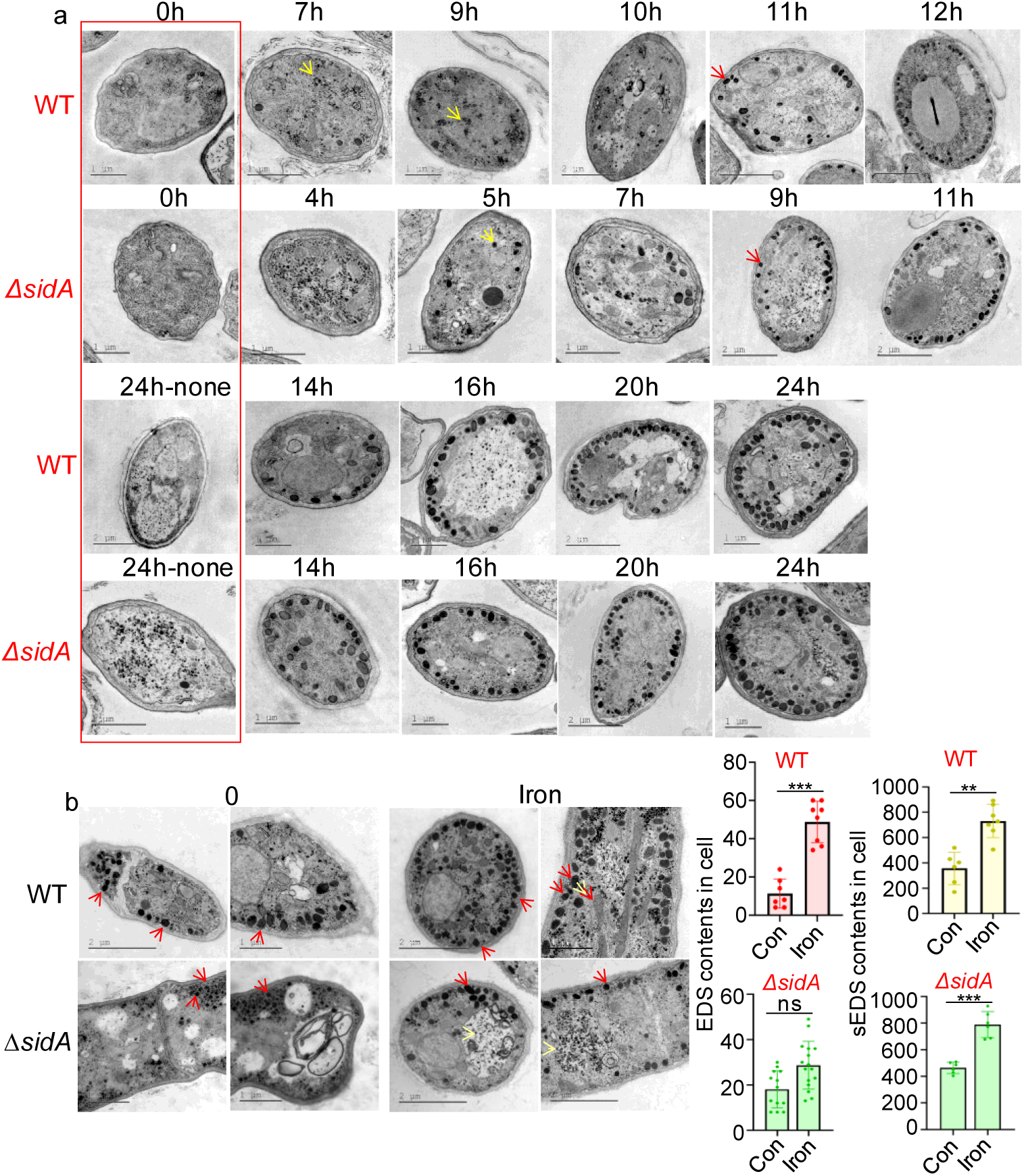
Evaluation of the role of iron in the contents of electron-dense bodies in trapping devices of *A. oligospora*. (**a**) TEM analysis of the formation of electron-dense bodies (EDSs) in WT and the mutant *ΔsidA* treated with nematodes within 24 h revealed that the lack of iron-chelating desferriferrichromes promoted the formation of electron dense bodies (EDSs) via accelerating the formation and aggregation of small EDSs (sEDSs) at early stage. Red arrows: EDS. Yellow arrows: sEDS. 24h-none: the strain without nematode treatment for 24h. (**b**) Comparison of the EDSs and sEDSs in trap cells of WT and the mutant *ΔsidA* treated with iron and solvent revealed that iron promoted the contents of both EDSs and (sEDSs in WT and the mutant *ΔsidA*. Significance was tested using an unpaired t test and adjusted using the Benjamini**−**Hochberg approach to control the false discovery rate (\**P* < 0.05, \*\**P* < 0.01, \*\*\**P* < 0.001, ns = not significant).

### 2. Iron enhanced the contents of electron dense bodies in trap cells

Iron was added to the culture of *A. oligospora* WT and the mutant Δ*sidA*, and the trapping devices of these strains were collected after treatment with the model nematodes for 24 hours. Both iron-treated strains developed significantly more trapping devices compared to their respective non-iron-treated counterparts. TEM analysis of the trap cells revealed that mature electron-dense bodies in both iron-treated strains were notably larger than those in the corresponding non-iron-treated strains (Fig. 2b). Specifically, the iron-treated Δ*sidA* exhibited larger, more well-defined, and tightly shaped mature electron-dense bodies than the non-iron-treated mutant Δ*sidA*. (Fig. 2b). Furthermore, the number of small electron-dense bodies in the trap cells of iron-treated WT was significantly higher compared to non-iron-treated WT (Fig. 2b). These results indicate that iron strongly enhances the formation and maturation of electron-dense bodies.

### 3. Iron is a key metal element in electron dense bodies

The elemental composition and distribution in trap cells of both *A. oligospora* WT and the mutant Δ*sidA* were evaluated using an energy-dispersive X-ray (EDX) detector integrated into a transmission electron microscopy (TEM) system, as described in a previous study [23]. Significant iron signatures were observed in the EDX spectra of electron-dense bodies and organelles, including mitochondria, vacuoles, and cell walls (Fig. 3a). Comparison of iron levels in these organelles within the same trap cells revealed that small electron-dense bodies contained significantly more iron than mature electron-dense bodies, mitochondria, vacuoles, and cell walls in both WT and Δ*sidA* trap cells (Figs. 3a–3b and Table S1). This iron enrichment in small electron-dense bodies was consistent with the increased iron content and the presence of small electron-dense bodies in the Δ*sidA* mutant, which lacks the iron-chelating desferriferrichromes. Remarkably, the iron content in small electron-dense bodies was more than twice as high as in mitochondria, the next largest iron reservoir (Fig. 3b and Table S1). Both small and mature electron-dense bodies were rich in iron, supporting the idea that iron promotes the formation of these bodies. Unexpectedly, vacuoles, which are typically the main iron storage organelles in fungi, contained the least iron in the trap cells of both WT and Δ*sidA* (Fig. 3 and Table S1).

**Fig 3.**
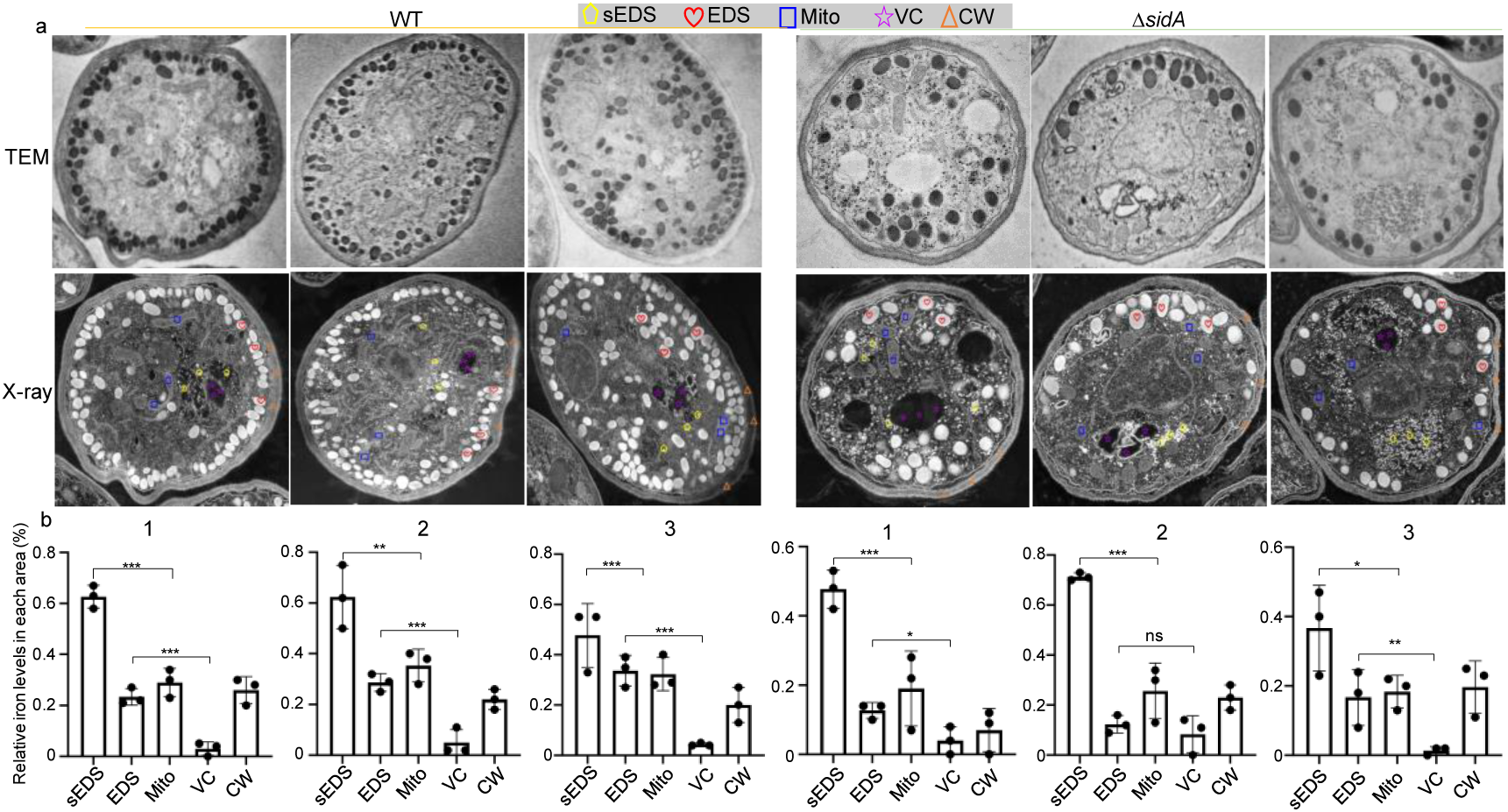
Evaluation of iron distribution and contents in trapping devices of *A. oligospora*. (**a**) TEM image and EDX electronic micrograph of trapping devices of WT and mutant Δ*sidA* with analysis sites marked by different color indicators. Yellow pentagon: sEDS; Red heart: EDS; Blue square: mitochondria (Mito); Purple five-pointed star: vacuole (VC); Orange triangle: cell wall (CW). (**b**) Comparison of relative iron contents in different organelles within the same trap cells of WT and mutant Δ*sidA* according to X-ray spectral analysis. Significance was tested using an unpaired t test and adjusted using the Benjamini**−**Hochberg approach to control the false discovery rate (\**P* < 0.05, \*\**P* < 0.01, \*\*\**P* < 0.001, ns = not significant).

### 4. Trapping devices harbor more iron than mycelia of *A. oligospora*

To assess the role of trapping devices in iron storage and isolation, potentially due to the inefficiency of vacuoles, we compared the iron levels between the mycelia and trapping devices of both *A. oligospora* WT and the mutant Δ*sidA*. Separating trapping devices from mycelia on a single plate is challenging, as the collected trapping devices typically contain substantial amounts of mycelia. Nevertheless, mixed samples of trapping devices and mycelia from both strains exhibited significantly higher levels of ferrous, ferric, and total iron compared to mycelia-only samples (Fig. 4a). Notably, the ferric iron levels in the mixed samples were 1.3-fold higher than those in the mycelia-only samples. Additionally, the mixed samples exhibited a markedly lower ratio of ferrous to ferric iron compared to the mycelia-only samples (Fig. 4a). These findings suggest that in *A. oligospora*, the trapping devices contain higher iron levels than the mycelia, consistent with the presence of iron-enriched electron-dense bodies in the trapping devices.

**Fig 4.**
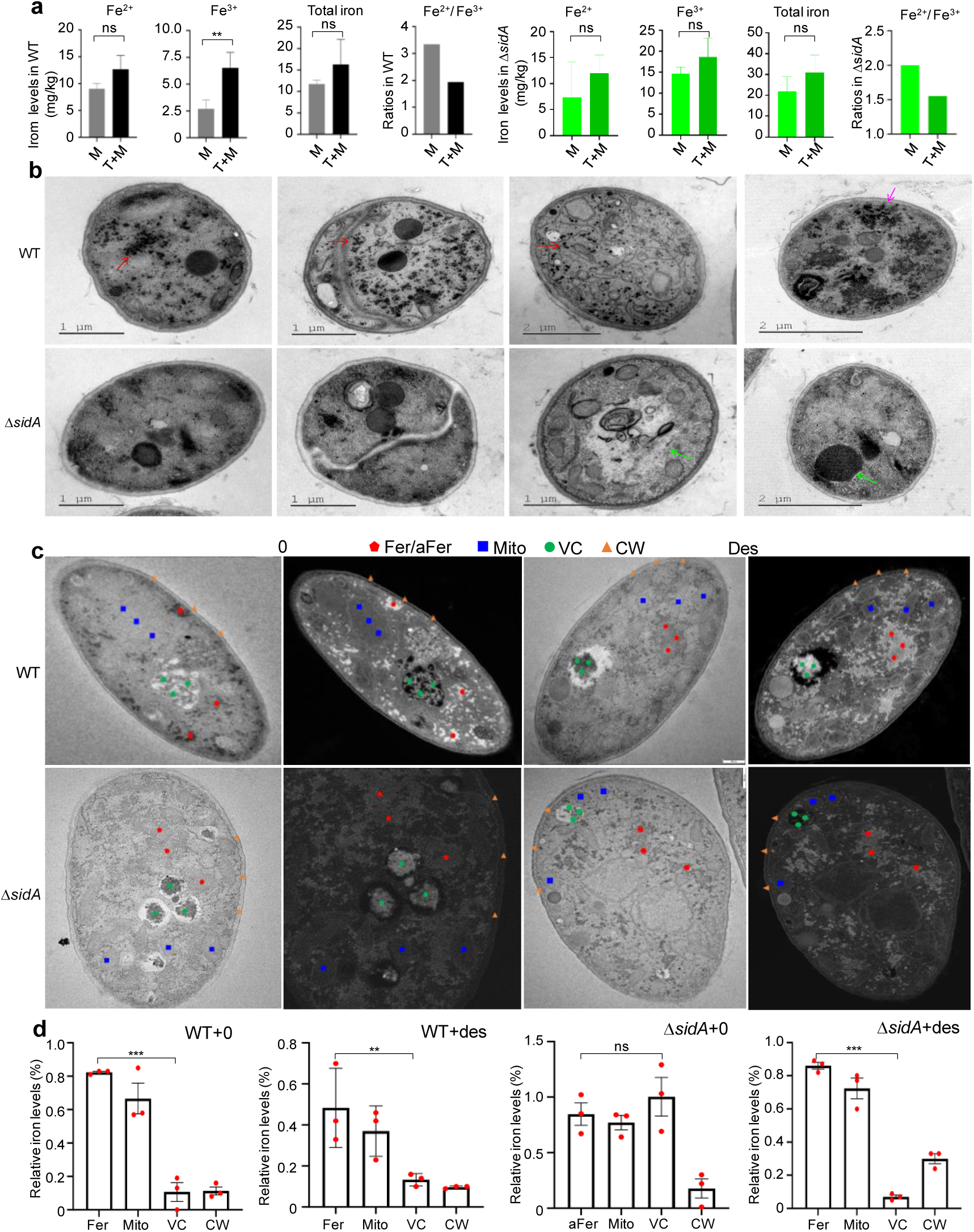
Evaluation of desferriferrichromes-mediated iron storage in the mycelia of *A. oligospora*. (**a**) Comparison of iron contents, including the levels of ferrous, ferric and total iron, and the ratios of ferrous to ferric iron, between trapping devices and mycelia in WT and the mutant Δ*sidA*. M: mycelia; M+T: mycelia and trapping devices. (**b**) TEM analysis of the mycelia cells between WT and the mutant Δ*sidA*. Red arrow: ferrichromes derived from desferriferrichromes; Purple arrow: aggregated ferrichromes. Green arrow: vacuoles (VC). (**c**) TEM-EDX electronic micrograph of the mycelia cells with analysis sites marked by different color indicators. Red pentagon: Ferrichrome (Fer) in WT while non-ferrichrome granules (aFer) in the mutant Δ*sidA* due to lack of desferriferrichromes. Blue square: mitochondria (Mito); Green dot: Vacuole (VC); Brown triangle: cell walls (CW). (**d**) Comparison of relative iron contents in various organelles within the same mycelia cell based on X-ray spectra analysis. Significance was tested using an unpaired t test and adjusted using the Benjamini**−**Hochberg approach to control the false discovery rate (\**P* < 0.05, \*\**P* < 0.01, \*\*\**P* < 0.001, ns = not significant).

### 5. Ferrichromes dominate iron storage in mycelia of *A. oligospora* rather than vacuoles

Fungal mycelia exhibit decision-making abilities, adjusting their developmental patterns in response to environmental changes by modulating metabolic and cellular processes [24]. We hypothesized that the increase in iron levels due to the absence of the iron-chelating desferriferrichromes in fungal mycelia might trigger mycelial differentiation into trapping devices for iron storage. To test this, we compared the mycelial cells of *A. oligospora* WT and the mutant Δ*sidA*. TEM analysis revealed that the mycelial cells of WT contained numerous ferrichrome granules, which were significantly reduced in the Δ*sidA* mutant (Fig. 4b). To further investigate the role of desferriferrichrome in ferrichrome granule formation, we supplemented the Δ*sidA* mutant with desferriferrichrome and used solvent as a control. Treatment with desferriferrichrome restored normal ferrichrome granule formation in the Δ*sidA* mutant mycelial cells, compared to solvent treatment, while no significant differences were observed in the WT treated with desferriferrichrome versus solvent alone (Fig. 4b). Notably, we also observed that vacuoles were present in the mycelial cells of the Δ*sidA* mutant but absent in WT mycelial cells (Fig. 4b).

Furthermore, desferriferrichrome treatment caused significant differences in the iron content of vacuoles in mycelial cells compared to solvent treatment, as revealed by EDX analysis (Figs. 4c–4d and Table S2). In the desferriferrichrome-treated Δ*sidA* mutant, vacuoles contained the least amount of iron among the organelles, whereas vacuoles in the solvent-treated mutant were the most iron-rich organelles (Figs. 4c–4d and Table S2). Notably, iron levels in ferrichrome granules were highest in the mycelia of strains containing desferriferrichromes, including the Δ*sidA* mutant treated with desferriferrichromes and WT strains treated with or without desferriferrichromes. This suggests that desferriferrichrome-mediated ferrichrome granules contain more iron than mitochondria and vacuoles (Figs. 4c–4d and Table S2). These results indicate that desferriferrichromes, rather than vacuoles, play a crucial role in iron storage in the mycelia of *A. oligospora*.

### 6. NTFs lack Ccc1-mediated vacuolar iron storage

To explore the genetic mechanisms underlying the non-effective vacuoles-mediated iron storage, we analyzed the distribution and abundance of 203 well-characterized vacuole-related genes of *Saccharomyces cerevisiae* in 9 NTFs and 13 non-NTFs genomes [15, 25]. Our analysis revealed that all NTFs lack 8 key vacuolar related genes (Fig. 5a), including, *Ccc1, thi4, ivy1, gnt1, vtc5, sna3, opy2* and *phm6*, which is consistent with the low iron storage observed in the vacuoles of *A. oligospora*. We further examined the distribution of the 8 vacuolar related genes across 2,057 high-quality fungal genomes (BUSCO complete scores > 90%) retrieved from the Mycocosm database. The results showed that *Ccc1* is highly conserved across the fungal kingdom (Fig. 5b and Table S3). A comparison of the prevalence of the *Ccc1* gene and the desferriferrichrome gene cluster in fungi revealed that 90.6% (1,864/2,057) of fungal genomes contain *Ccc1*, while only 45% (927/2,057) contain the desferriferrichrome gene cluster (Fig. 5b and Table S3). The proportion of genomes possessing both the *Ccc1* gene and the desferriferrichrome gene cluster was 40.7% (838/2,057). These results suggest that Ccc1-mediated vacuolar iron storage plays a crucial role in maintaining iron homeostasis in most fungi, but not in NTFs.

**Fig 5.**
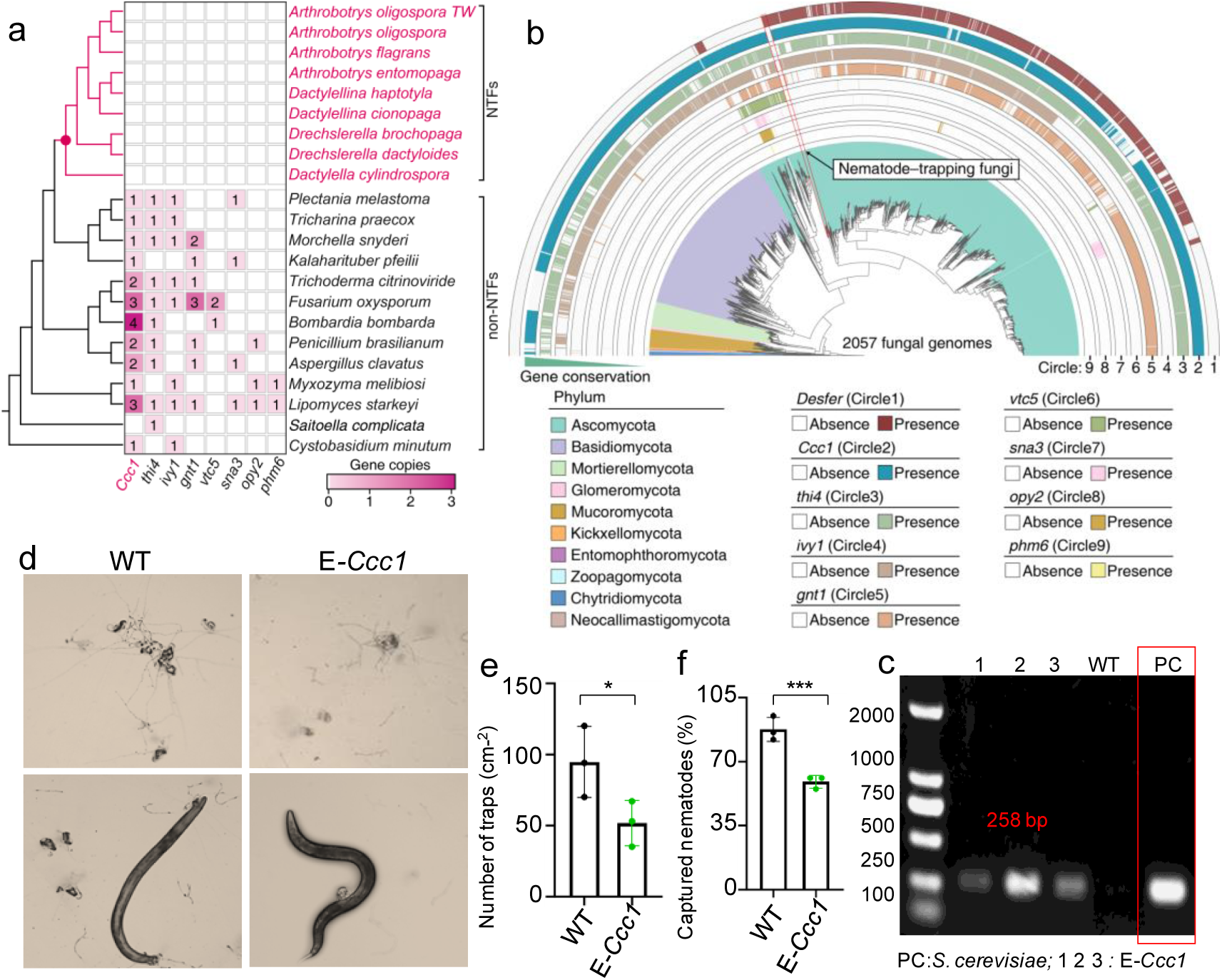
Phylogeny of vacuolar-related genes in Ascomycota and distribution of the *Ccc1* gene and the desferriferrichrome gene cluster in fungal group. (**a**) Heatmap illustrating the loss of eight genes involved in vacuolar related genes in NTFs, including vacuolar iron transporter (Ccc1), Thiamine thiazole synthase (thi4), Protein IVY1 (ivy1), lucose N-acetyltransferase 1 (gnt1), Vacuole transporter chaperone complex subunit 5 (vtc5), Protein SNA3 (sna3), Overproduction-induced pheromone-resistant protein 2 (opy2) and Phosphate metabolism protein 6 (phm6). The copy numbers of each gene are indicated in each color block. The tree topological structure is derived from the 2075 fungal genome tree. (**b**) Distribution patterns of the desferriferrichrome gene cluster and vacuolar-related genes lost in NTFs in fungi kingdom. This panel highlights the presence and absence of these genes in fungal genomes. (**c**) Transcriptional analysis the exogenous gene *Ccc1*of *A. oligospora* mutant E-*Ccc1*with harboring the *Ccc1* gene from yeast. (**d−f**) Trap induction and nematicidal activities of WT and the mutant E-*Ccc1* with harboring an exogenous *Ccc1* gene from yeast. Significance was tested using an unpaired t test (\**P* < 0.05, \*\**P* < 0.01, \*\*\**P* < 0.001, ns = not significant).

Further analysis of the ferritin gene, which encodes a ubiquitous protein responsible for iron storage in both prokaryotes and eukaryotes [26–28], revealed variability in its copy numbers among NTFs, dependent on the type of trapping devices present. Specifically, one copy was found in species with adhesive networks (*Arthrobotrys oligospora* and *Arthrobotrys flagrans*), two copies in species with constricting rings (*Drechslerella brochopaga* and *Drechslerella dactyloides*), three copies in species with adhesive columns (*Dactylellina cionopaga*), and four copies in species with adhesive knobs (*Dactylellina haptotyla* and *Dactylellina entomopaga*) (Fig. S2a). Notably, the transcriptional level of the ferritin gene in *A. oligospora* is very low (average TPM = 4.19) and remains unaffected by the absence of desferriferrichromes (Fig. S2b). Moreover, NTFs lack the ability to harbor ferrosomes, a class of iron storage organelles encoded by the *fezABC* operon found in various bacteria and archaea, such as *Desulfovibrio magneticus*, due to the absence of homologous genes *fezA* and *fezC* [23, 29]. Furthermore, the transcriptional levels of most *fezB* genes showed no significant change between the Δ*sidA* and WT strains (Fig. S2c).

To verify the effect of the lack of *Ccc1* gene on the formation of trapping devices, a 969-bp fragment of the *Ccc1* gene was directly cloned from *Saccharomyces cerevisiae* BJ5464 and inserted into *A. oligospora* WT to generate a mutant E-*Ccc1* (Fig. S3). Transcriptional analysis confirmed that the *Ccc1* gene was successfully expressed in the E-*Ccc1* strain (Fig. 5c). Phenotypic analysis revealed that the E-*Ccc1* mutant exhibited a significant reduction in both trapping device formation and nematicidal activity compared to WT (Fig. 5d–5f).

### 7. The origin of trapping devices is related to heat-induced iron storage

To elucidate the evolutionary trajectory of NTFs, we identified 1,136 sets of 1:1 orthologous protein-coding genes across 22 fungal genomes described above and reconstructed a well-supported genome-wide phylogeny based on 1,141,520 concatenated amino acid positions (Fig. 6). A time-calibrated phylogenetic tree was constructed using 275,133 four-fold degenerate sites from 1,136 single-copy orthologous genes, calibrated with three major events [30]: the Ascomycota crown group (487–773 Mya), Orbiliomycetes–other Pezizomycotina (353–554 Mya), Saccharomycotina crown group (276–514 Mya), and one fossil calibration point (A extinct NTF group with ring trapping structure: >100 Mya) [31]. The resulting time-calibrated phylogenetic tree revealed that the last common ancestor (LCA) of NTFs in the Ascomycota existed around 255 Mya (CI: 231–287 Mya), which is consistent with previous studies suggesting that the origin of trapping devices in NTFs coincided with the Permian-Triassic mass extinction.

**Fig 6.**
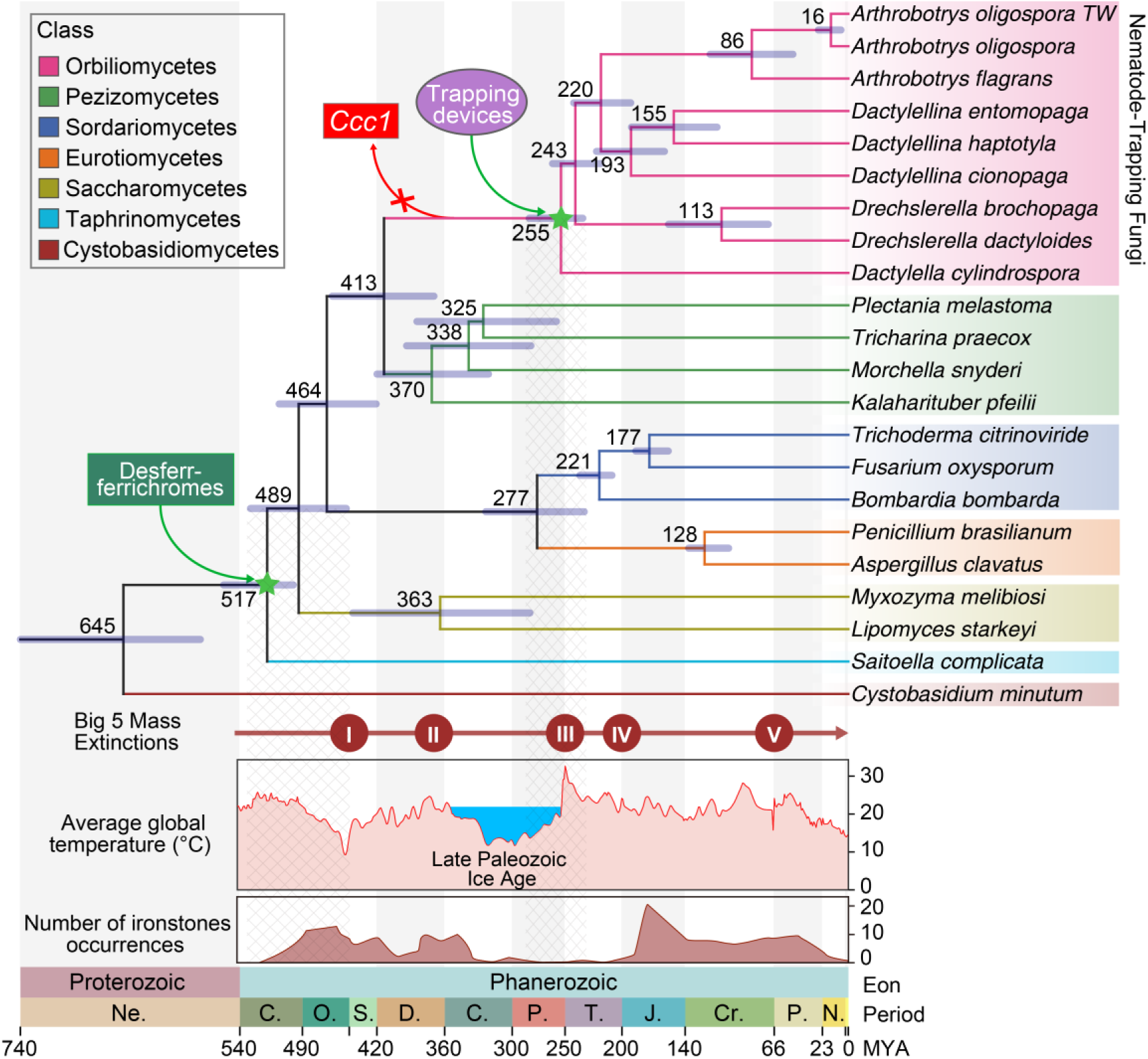
Estimated divergence times of major lineages in the carnivorous Orbiliomycetes, with inferred occurrences of Phanerozoic ironstones and Phanerozoic global average temperature (GAT) trends. The chronogram was constructed using MCMCTree in the PAML package with 275,133 four-fold degenerate sites from 1,136 single-copy orthologous genes and calibrated with three major events: Ascomycota crown group (487–773 Mya), Orbiliomycetes–other Pezizomycotina (353–554 Mya), Saccharomycotina crown group (276–514 Mya) and one fossil calibration point (A extinct NTF group with ring trapping structure: >100 Mya). Error bars represent the 95% highest posterior density (HPD) of node ages. Numbered circles at each node indicate node sequence from root to leaves, and node ages are shown. Circles labeled with Roman numerals represent major Phanerozoic mass extinction events: Late Devonian (375 Mya), Permian-Triassic (251.4 Mya), Triassic-Jurassic (201.4 Mya), and Cretaceous-Paleogene (66 Mya). Time periods are abbreviated as follows: De: Devonian, Ca: Carboniferous, Pe: Permian, Tr: Triassic, Ju: Jurassic, Cr: Cretaceous, P: Paleogene, N: Neogene. The gain and loss of key genes and the appearance of trapping device were marked on the corresponding tree nodes.

Mapping iron content fluctuations over geological time throughout the Phanerozoic onto the evolutionary trajectory of carnivorous fungi reveals that the origin of the fungal lineage containing the desferriferrichromes gene cluster occurred around 517 Mya (CI: 494–555 Mya), aligning with a significant increase in ironstone during the early Ordovician (Fig. 6). However, no correlation between geological iron content changes and the Permian-Triassic mass extinction and the origin of trapping devices in NTFs was observed. Many studies suggest that the Permian-Triassic mass extinction is linked to a sharp rise in global temperatures, with sea surface temperatures reaching approximately 40°C and land temperatures likely fluctuating even higher [32–35]. When the global average temperatures over the Phanerozoic are mapped onto the evolutionary trajectory of carnivorous fungi, the origin of trapping devices coincides with lethally high temperatures during the early Triassic Greenhouse (Fig. 6).

Interestingly, the emergence of the desferriferrichromes gene cluster also coincided with a period of high temperatures during the Cambrian (Fig. 6), suggesting a link between desferriferrichromes-mediated iron storage and high temperature. Notably, the ancestors of NTFs endured the longest ice age, the Late Paleozoic Ice Age (LPIA), from 360 to 255 Mya, after which NTFs with trapping devices emerged during the intense heat of the early Triassic Greenhouse [36, 37]. Importantly, the crucial vacuolar gene *Ccc1* was lost during the Late Paleozoic Ice Age (Fig. 6). The loss of Ccc1-mediated vacuolar iron storage under low temperatures aligns with the gain of desferriferrichromes-mediated iron storage under high temperatures, suggesting that high temperatures promote iron storage, whereas low temperatures have the opposite effect. Thus, the emergence of trapping devices in NTFs during the extreme heat following a long period of low temperatures may represent a new strategy for iron storage induced by high temperature.

### 8. Heat promotes trap-mediated iron storage

We assessed the effects of a temperature gradient (from 32°C to 12°C) on the growth and trapping device formation of *A. oligospora* WT and the mutant Δ*sidA* on PDA at six different temperatures: 32°C, 28°C, 24°C, 20°C, 16°C, and 12°C. Notably, both WT and Δ*sidA* colonies exhibited the largest growth at 24°C, followed by the second-largest growth at 28°C and 20°C, the third-largest growth at 16°C, and the smallest growth at 12°C (Fig. 7a). Importantly, both strains failed to grow at 32°C but successfully grew at 12°C, indicating that *A. oligospora* is sensitive to high temperatures. Under nematode treatment, both WT and Δ*sidA* formed the most trapping devices at 28°C, followed by the second-highest number at 24°C, the third-highest at 20°C (Figs. 7b and 7c), and the fewest at 12°C, demonstrating a temperature-dependent formation of trapping devices. For the Δ*sidA* mutant, trapping device formation increased progressively as the temperature rose from 12°C to 28°C (Figs. 7b and 7c). Additionally, both strains displayed temperature-dependent nematicidal activity across the gradient from 28°C to 12°C (Figs. 7b and 7c).

**Fig 7.**
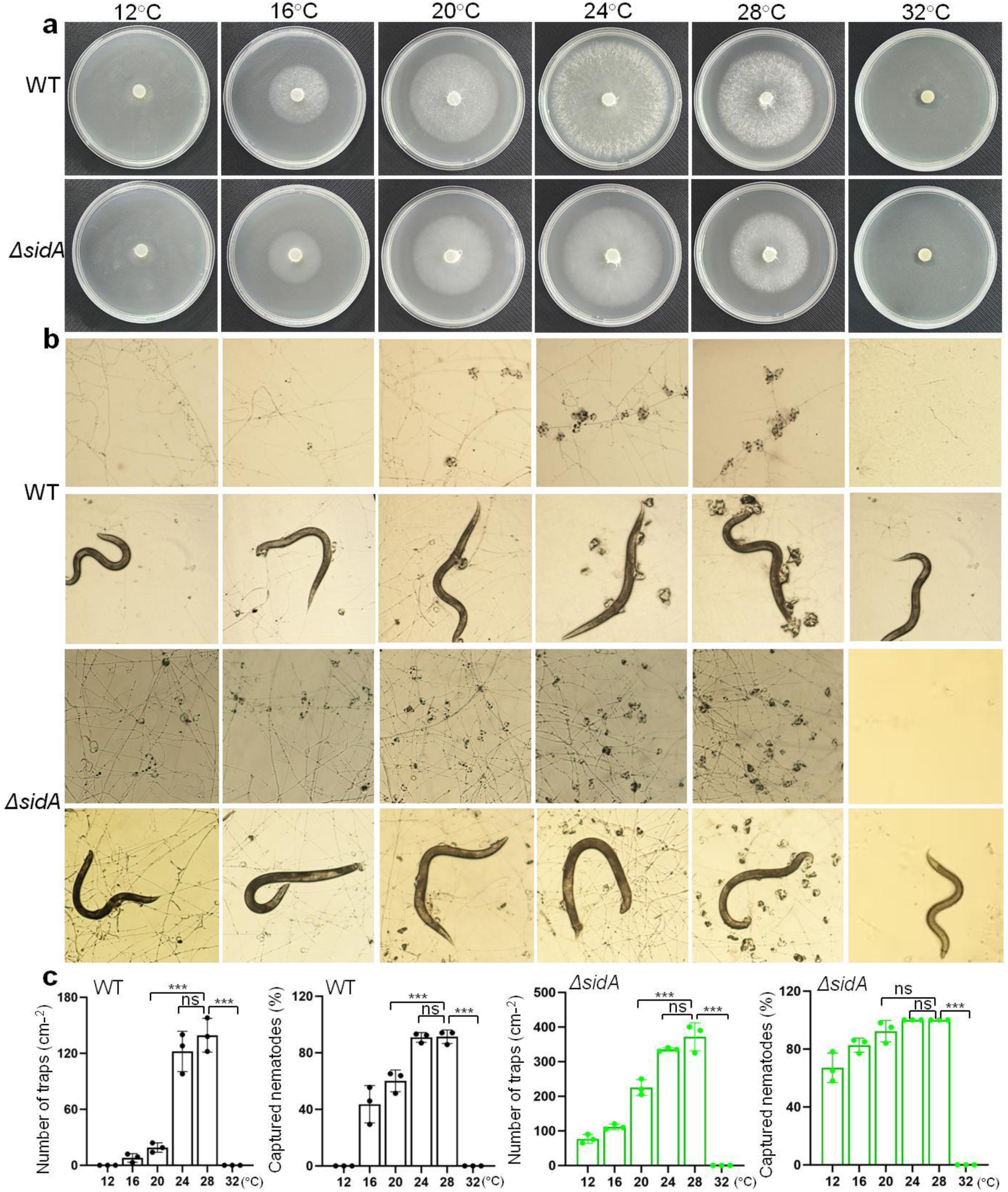
Evaluation of the effect of temperatures on colony growth, trapping devices, and nematicidal activity in *A. oligospora*. (**a**) Pictures of colony growths of WT and the mutant Δ*sidA* at different temperatures of 32°C, 28°C, 24°C, 20°C, 16°C and 12°C. (**b**) Pictures of trapping devices and captured nematodes of WT and the mutant Δ*sidA* at 32°C, 28°C, 24°C, 20°C, 16°C and 12°C. (**c**) Comparison of trapping devices and captured nematodes of WT and the mutant Δ*sidA* at 32°C, 28°C, 24°C, 20°C, 16°C and 12°C. Significance was tested using an unpaired t test and adjusted using the Benjamini**−**Hochberg approach to control the false discovery rate (\**P* < 0.05, \*\**P* < 0.01, \*\*\**P* < 0.001, ns = not significant).

Previous studies have shown that water agar medium is widely used to induce the formation of trapping devices in *A. oligospora* without the addition of nematodes. To further investigate the effect of temperature on trapping device formation, we cultivated both WT and the Δ*sidA* mutant on water agar at three temperatures: 28°C, 20°C, and 12°C. As expected, both WT and Δ*sidA* strains spontaneously developed numerous trapping devices at 28°C, while the WT strain failed to form trapping devices at 12°C (Fig. 8a). Both strains exhibited the largest and most abundant trapping devices at 28°C, followed by fewer and smaller trapping devices at 20°C, and the least and smallest at 12°C (Fig. 8a). These results confirm that trapping device formation in *A. oligospora* is significantly temperature-dependent.

**Fig 8.**
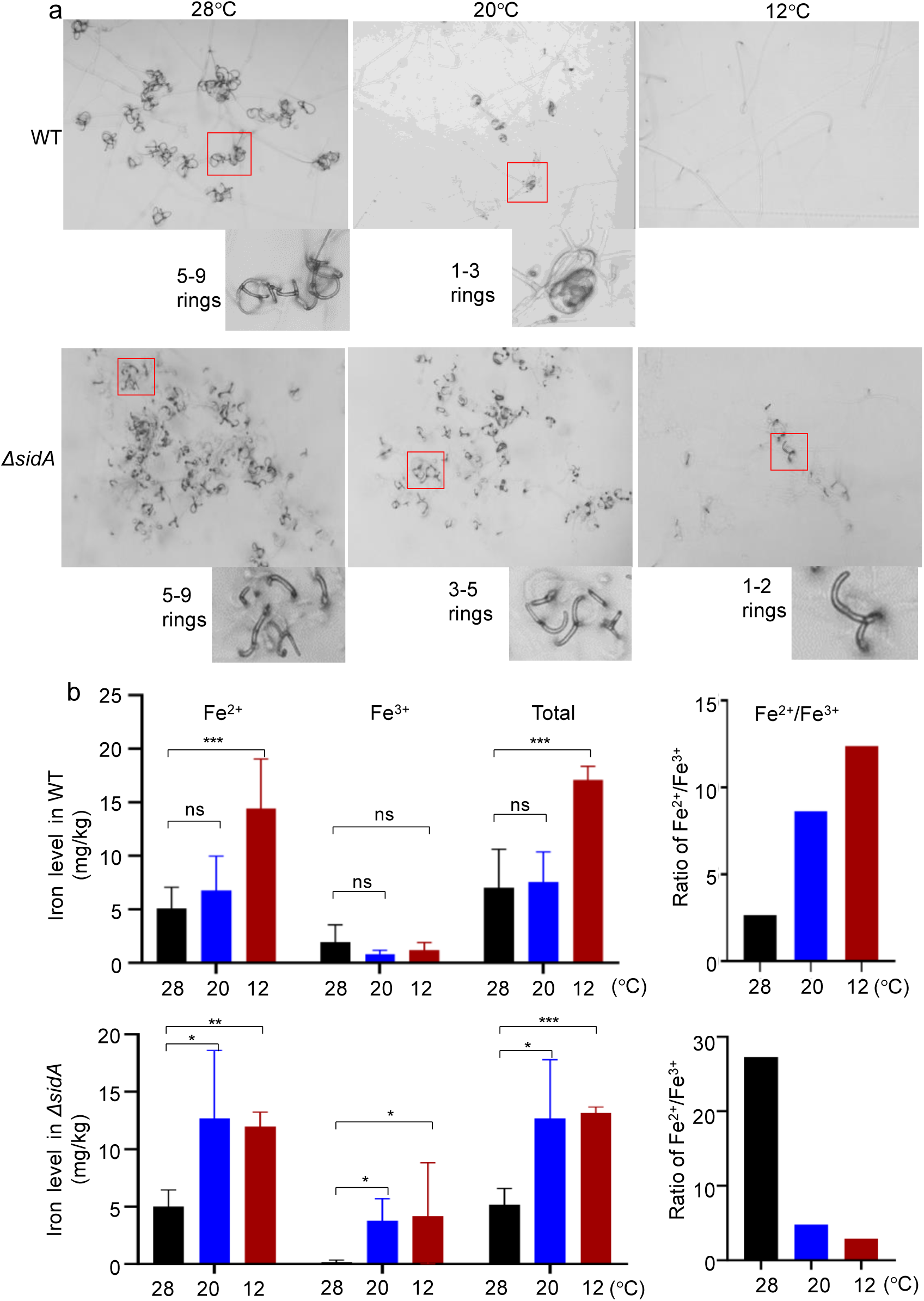
Evaluation of temperatures-mediated relationship between trapping devices and iron contents in *A. oligospora*. (**a**) Pictures of trapping devices of WT and the mutant Δ*sidA* at 28°C and 20°C, and 12°C. (**b**) Comparison of iron contents, including the levels of ferrous, ferric and total iron, and the ratios of ferrous to ferric iron, in WT and the mutant Δ*sidA* at 28°C and 20°C, and 12°C. Significance was tested using an unpaired t test and adjusted using the Benjamini**−**Hochberg approach to control the false discovery rate (\**P* < 0.05, \*\**P* < 0.01, \*\*\**P* < 0.001, ns = not significant).

**Figure 9.**
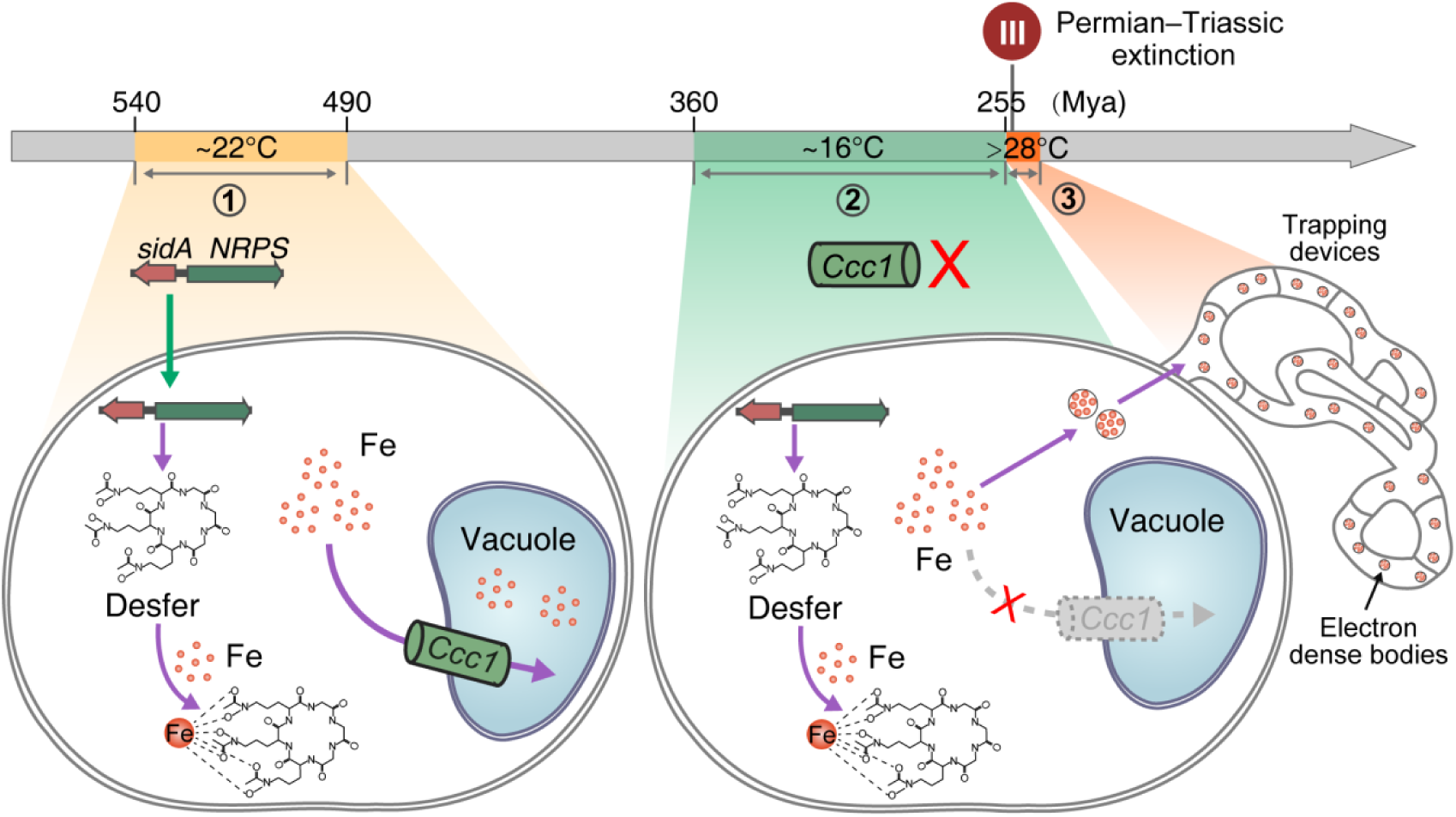
Diagram of the roles of temperature-mediated iron storage in the origin and evolution of trapping devices in NTFs. The desferriferrichromes gene cluster emerged in fungi during the high temperatures in the Cambrian (540−490Mya). The ancestors of NTFs lost Ccc1-mediated vacuolar iron storage during the longest ice age of the Late Paleozoic Ice Age (360−255 Mya). The emergence of trapping devices in NTFs occurred during the deadly heat after the longest ice age.

Previous studies have shown that the free iron content in fungi is negatively correlated with temperature [38, 39]. For example, thermophilic fungi require higher levels of free iron to facilitate the Fenton reaction, which generates energy for adaptation to relatively low ambient temperatures. To investigate the effect of temperature on free iron content in *A. oligospora*, we compared the iron levels and ratios in the mycelia of both WT and Δ*sidA* at three temperatures: 28°C, 20°C, and 12°C. We observed that the WT strain had the lowest ferrous and total iron contents at 28°C and the highest at 12°C (Fig. 8b). Similarly, the Δ*sidA* mutant showed the lowest ferric and total iron contents at 28°C and the highest at 12°C (Fig. 8b). These results support the notion that free iron content in fungi is inversely correlated with temperature, suggesting that NTFs exposed to relatively high temperatures must store iron to reduce its availability in the mycelia.

Notably, WT exhibited the lowest ratio of ferrous to ferric iron at 28°C and the highest at 12°C (Fig. 8b), indicating a negative correlation between the ferrous-to-ferric ratio and temperature. In contrast, the Δ*sidA* mutant showed the highest ratio of ferrous to ferric iron at 28°C and the lowest at 12°C (Fig. 8b). Excess intracellular ferrous iron can be highly toxic, as misregulated levels of this reactive metal increase the susceptibility of organisms to harmful Fenton-like reactions, which produce highly reactive hydroxyl radicals. The absence of the iron-chelating desferriferrichromes in the Δ*sidA* mutant likely caused the reversal of the ferrous-to-ferric ratio-temperature relationship observed in WT. Therefore, desferriferrichromes play a crucial role in maintaining the negative correlation between the ferrous-to-ferric ratio and temperature in *A. oligospora*, thereby protecting against excess ferrous ions.

## Discussion

The ability to segregate superfluous ingredients in response to environmental changes is essential for survival and fitness in all organisms. Temperature is a critical environmental and climatic selective force that influences morphological changes by affecting essential physiological and biological processes [40–43]. Under some conditions, mycelia produce spores either directly or through specialized fruiting bodies. Fungal mycelia can be microscopic or develop into visible structures, such as brackets, mushrooms, puffballs, rhizomorphs (long strands of hyphae cemented together), sclerotia (hard, compact masses), stinkhorns, toadstools, and truffles [44]. Recent phylogenetic analyses of the desferriferrichrome biosynthetic enzymes suggest that NTFs are closely related to species such as *Kalaharituber pfeilii* (a desert truffle), *Tricharina praecox* (cup fungus), *Trichophaea hybida* (cup fungus), *Morchella sextelata* (true morel), *Morchella importuna* (true morel), *Choiromyces venosus* (a truffle), and *Tuber borchii* (a truffle) [14]. Many of these fungi produce valuable and edible fruiting bodies. NTFs develop trapping devices from vegetative mycelia as well. The formation of trapping devices in NTFs is considered a diagnostic marker for the transition from a saprophytic to a predatory lifestyle and plays a crucial role in their carnivorous activity. While nutrient signals have historically been regarded as the primary triggers for trapping device formation [21], studies on the role of temperature in this process remain largely unexplored.

In this study, we investigated the relationship between the origin of trapping devices and iron levels throughout the Phanerozoic. Unexpectedly, we found that the origin of trapping devices was not linked to changes in iron levels, but rather to temperature fluctuations. The emergence of trapping devices coincided with the extreme heat of the early Triassic Greenhouse, following the longest ice age, the Late Paleozoic Ice Age. Importantly, the loss and acquisition of key genes responsible for iron storage were also strongly associated with temperature changes. The Ccc1-mediated vacuolar iron storage mechanism was lost during the Late Paleozoic Ice Age, while the desferriferrichrome-mediated iron storage pathway was acquired during the high temperatures of the Cambrian period (Fig. 6).

Phenotypic analysis revealed that the formation of trapping devices in *A. oligospora* is positively correlated with temperature, indicating that temperature is an abiotic factor that triggers trapping device formation in NTFs. Additionally, we observed a negative correlation between temperature and iron content in *A. oligospora*, with mycelia grown at high temperatures exhibiting lower iron levels. Importantly, we found that the trapping devices in *A. oligospora* contained higher iron levels than the mycelia. TEM-EDS analysis of iron distribution and content in trap cells showed that the electron-dense bodies in the trapping devices were enriched with iron. Furthermore, supplementation with the iron chelator desferriferrichrome significantly inhibited trapping device development in *A. oligospora*. Conversely, iron supplementation or a deficiency in desferriferrichrome biosynthesis triggered extensive trapping device formation.

Notably, TEM-EDS analysis revealed that the electron-dense bodies within these trapping devices are more efficient than mitochondria and vacuoles in iron storage, although they are less effective than desferriferrichrome-mediated ferrichrome granules in mycelia. These electron-dense bodies serve as an alternative iron storage system. The results suggest that the heat-sensitive *A. oligospora* has developed trapping devices to sequester excess iron and reduce iron content in mycelia when exposed to heat, a key adaptive strategy for responding to temperature fluctuations. A previous study reported that electron-dense bodies in the alga *Cyanidium caldarium* also function in iron storage [45]. This suggests that the role of electron-dense bodies in iron sequestration may be a widespread phenomenon, rather than specific to a single fungal strain. The key difference between *A. oligospora* and *C. caldarium* is that the fungus develops specialized trapping devices to store these iron-rich electron-dense bodies. We propose that electron-dense bodies in *C. caldarium* may also play a role in the algal adaptation to temperature changes, which warrants further investigation.

The ability to temporarily store excess components to adapt to environmental changes is a fundamental aspect of homeostasis regulation in all living organisms. Iron overload can lead to cellular toxicity, primarily through the generation of reactive oxygen species (ROS) and lipid membrane peroxidation [46]. Hemochromatosis is a disorder characterized by iron overload resulting from both inherited and acquired factors [47, 48]. If left untreated, excess iron can cause damage to various organs, including the liver, heart, pancreas, endocrine glands, and joints, potentially leading to cirrhosis, liver failure, cancer, arthritis, diabetes, and heart failure [47]. Caucasians are particularly susceptible to developing iron overload and the associated complications of hemochromatosis due to a higher prevalence of mutations in the homeostatic iron-regulator gene within this population, which leads to excessive iron absorption [49, 50]. Here, we show that heat is also an environmental factor contributing to iron excess, which provides a new mechanism for controlling iron availability, and mitigating its intrinsic toxicity to improve treatments for iron overload.

In summary, we have identified that the iron-rich electron-dense bodies in trapping devices represent a novel type of microbial iron-storage granules. Remarkably, the trapping devices serve as a unique iron storage system, distinct from the iron-storing granules or organelles previously reported in cells. Our results suggest that *A. oligospora* could serve as a potential eukaryotic model with an iron overload phenotype, which may be applied to develop innovative agents and therapies for treating iron overload-associated diseases. In this model, temperature is identified as the first non-nutrient factor that triggers the development of trapping devices diagnostic for iron overload in *A. oligospora*. Our results suggested that NTFs could serve as a potential eukaryotic model for elucidating dynamic iron homeostasis regulation, which may aid in the treatment of iron overload-related diseases.

## Acknowledgment

We would like to thank the School of Chemical Science and Technology, Key Laboratory of Medicinal Chemistry for Natural Resource - Ministry of Education, Yunnan University for providing us with TEM-EDX. We wish to thank Longchun Bian, for support with the Energy Dispersive X-ray Spectroscopy analyses and data reduction. We would like to thank the Institutional Center for Shared Technologies and Facilities of Kunming Institute of Zoology, Chinese Academy of Sciences, for providing us with TEM. We are grateful to Yingqi Guo for her great help in preparing TEM samples and taking/analyzing TEM images.

## Data availability

Our sequences files of transcriptome are accessible from the National Center for Biotechnology Information (NCBI) under BioProject accession: PRJNA639935 (https://www.ncbi.nlm.nih.gov/bioproject/PRJNA1073725).

