## supporting information for "Nematode-Trapping Devices of *Arthrobotrys oligospora* is an Iron Storage Phenotype Adapted to Temperature Changes"

**Materials and Methods**

**Fungal strains and cultivation**

YMF1.01883 (ATCC 24927) was procured from the State Key Laboratory for Conservation and Utilization of Bio-Resources and was cultivated at 25 °C for 3 to 4 days on a 9 cm Petri dish containing PDA (potato 200 g/L, glucose 10 g/L, agar 15 g/L, ddH_2_O or tap water). Subsequently, 9 mm discs of the fungal colonies were transferred to 9 cm Petri dishes containing YMA (2 g/L yeast extract, 10 g/L maltose, agar 20 g/L) and incubated at 25 °C for 6 to 7 days. The spores were then washed with 20 mL of 0.05% Tween-80 mixed with approximately 3 mm glass beads.

**Generation of gene deletion mutants**

A modified homologous recombination method was employed to construct the mutants Δ*SidA* and Δ*NRPS*. The upstream and downstream homologous arms of the target gene were cloned from *A. oligospora* YMF1.01883, and the *HygR* resistance sequence was cloned from a PUC19-1300-D-HYB vector using the Prime STAR Max enzyme (TsingKe, Beijing, China). All primers used in this study are listed in Table S4. The DNA fragments (5′ flank, hygromycin B, and 3′ flank) were purified using the RiboEXTRACT Universal DNA Purification Kit (Tsingke, China). The purified 5′ flank DNA fragment was inserted into the enzyme-digested PUC19-1300-D-HYB vector using the In-Fusion method to produce a PUC19-1300-D-HYB-5′ vector. Subsequently, the 3′ flank DNA fragment was inserted into this vector to generate a PUC19-1300-D-HYB-5′-3′ vector. The homologous fragment was amplified and purified as previously described. *A. oligospora* protoplasts were prepared as previously described. The purified homologous fragment was transformed into *A. oligospora* protoplasts. Transformation colonies were selected after incubation at 28 °C for 2−4 days, and each colony was transferred to a new plate containing PDA. After incubation at 28 °C for 5 days, genomic DNA of putative transformants was extracted and verified by polymerase chain reaction (PCR) to check for the integration of the target gene into the genome. Mutants deficient in the target gene were screened by PCR and confirmed by sequencing analysis. Further confirmation of the mutants was performed by Southern blot hybridization using the North2South Chemiluminescent Hybridization and Detection Kit according to the manufacturer’s instructions (Pierce, Rockford, IL). For Southern blot analysis, restriction enzyme HindIII was used to digest WT and Δ*SidA* genomic DNA, and restriction enzyme SmaI was used to digest WT and Δ*NRPS* genomic DNA (Fig S3).

**Construction of mutant E-*Ccc1***

To construct a heterologous expression plasmid, the PUC19-1300-D-HYB plasmid was amplified by means of fusion polymerase chain reaction (PCR) using Phanta Max SuperFidelity DNA Polymerase (Vazyme, Nanjing, China) according to the manufacturer’s protocol. All primers used in this study are provided in Table S5. The heterologous fragment was amplified from *Saccharomyces cerevisiae* BJ5464 and purified as described previously. Gene *Ccc1* was inserted into PUC19 plasmid containing *PtrpC* and *TtrpC* using strong promoter *PtrpC* and terminator *TtrpC* from *Aspergillus nesterus*. The plasmid was transformed into *A. oligospora* protoplasts respectively. Transformation colonies were selected after incubation at 28 °C for 2−4 days, and every single colony was transferred to a new plate containing PDA. After incubation at 28 °C for 5 days, genomic DNA of putative transformants was extracted as described previously and verified by PCR to check for integration of the target in the genome. The mutant was screened out and confirmed by PCR and sequencing analysis. The WT strain and E-*Ccc1* mutant were cultured on PDA medium at 28°C for 6 days. The vegetative hyphae were harvested in 9-cm Petri dishes, which were then incubated at 28 °C for 3 and 5 d. After induction, hyphae were collected and frozen immediately in liquid nitrogen. Total RNA was extracted from all samples with the AxyPrep Multisource RNA Miniprep Kit (Axygen, Jiangsu, China). The extracted RNA was then reverse transcribed to cDNA with the FastQuant RT Kit with gDNase (Takara, Kusatsu, Japan). The cDNA was used as the template for analysis of expression of gene *Ccc1.* The cDNA of *Saccharomyces cerevisiae* BJ5464 was used as the template for positive control.

**Development of trapping device**

To facilitate the collection of mycelia, *A. oligospora* YMF1.3170 was cultured at 25 °C for 3 to 4 days on 9 cm Petri dishes containing potato dextrose agar (PDA) with cellophane (4 cm diameter, area 50.24 cm²). *Caenorhabditis elegans* N2 was grown in a 250 mL flask with oatmeal agar medium (oatmeal 30 g, water 20 mL) for 7 days. The nematodes were then collected from the flask's walls and transferred to a 2 mL microcentrifuge tube [51]. To prepare the nematode suspension, 1 mL of double-distilled water was added and mixed. The nematode concentration was subsequently adjusted to 300 nematodes per 200 µL for the assay. The wild-type (WT) and mutant strains were exposed to 200 µL of double-distilled water containing 300 nematodes to induce the formation of trapping devices, classified as the mycelia + trapping devices (M+T) group. The control group, which only included mycelia (M), was treated with 200 µL of double-distilled water without nematodes.

**Desferriferrichrome assay**

Five milligrams of desferriferrichrome (CAS No. 34787-28-5, Sigma Co.) were dissolved in 5 mL of dimethyl sulfoxide (DMSO) to prepare a 1 mg/mL desferriferrichrome stock solution. To achieve a final concentration of 1.45 μM desferriferrichrome, 150 μL of the 1 mg/mL stock solution was added to 150 mL of potato dextrose broth (PDB). Wild-type (WT) and mutant strains, cultured on potato dextrose agar (PDA) for 7 days, were inoculated into potato dextrose (PD) broth (200 g/L potatoes, 20 g/L glucose) and incubated on a shaker at 180 rpm for 12 hours. The culture was then supplemented with 1.45 μM desferriferrichrome and further incubated on the shaker for another 12 hours. The mycelia were harvested, fixed with 2.5% glutaraldehyde, and examined using transmission electron microscopy (TEM).

**Fe^3+^Assay**

The fungal strains were initiated with 9 cm diameter hyphal disks at 28℃ for 6 days on the plates of PDA supplemented with or without FeCl_3_·6H_2_O (10−30 μM) and then about 400 nematodes were introduced to the cultures. After 12 h, traps and captured nematodes per plate were observed under a microscope and counted at specific time-points. The diameter of each colony was measured and then calculated with GraphPad 9.3.4.

**Iron contents**

An iron assay kit (#ab83366, Abcam) was used to evaluate intracellular iron levels (Fe²⁺ and Fe³⁺) in fungi [38]. First, mycelia (20–80 mg) were collected in phosphate-buffered saline (PBS) containing 80 g NaCl, 2 g KCl, 14.4 g Na₂HPO₄, and 2.4 g KH₂PO₄ in 1 L of double-distilled water. The mycelia were then homogenized in iron assay buffer using a Dounce homogenizer on ice with 50–100 passes. Next, iron reducer was added to the collected supernatant, mixed, and incubated. Following this, iron probe (3-(2-Pyridyl)-5,6-bis(5-sulfo-2-furyl)-1,2,4-triazine, disodium salt) was added, mixed, and incubated for 1 hour. The resulting solution was immediately measured using a microplate reader at OD = 593 nm. To account for the effects of nematodes on iron content in the mycelia + trapping devices (M+T) groups, nematodes were removed from the M+T samples using a 10 µL pipette. Additionally, the iron content of 300 nematodes was measured separately. Iron content in the M+T groups, both with and without nematodes, was evaluated using the same method. All mycelia were collected, weighed, and suspended in double-distilled water at a 1:10 ratio. The suspension was ground using an ultrasonic cell disruptor to obtain a uniform mixture, from which 1 mL was taken for analysis. Similarly, 300 nematodes were suspended in 1 mL of distilled water, ground using an ultrasonic cell disruptor, and then centrifuged at 16,000 rpm for 10 minutes to separate the supernatant for further experiments.

**Transmission electron microscopy (TEM)**

For transmission electron microscopy (TEM) analysis, ultrathin sections of hyphal cells were prepared and examined as previously described [38]. The samples were pretreated with 4% paraformaldehyde and then incubated overnight at 4 °C with 2.5% glutaraldehyde (Sigma) in phosphate buffer (pH 7.4). Following this, the samples were post-fixed in 1% osmium tetroxide (OsO₄). They were then dehydrated through a gradient ethanol series, embedded in Spurr resin, and stained with 2% uranyl acetate and Reynold’s lead citrate. Finally, the samples were examined using an H-7650 transmission electron microscope (Hitachi, Tokyo, Japan).

**Transmission electron microscopy-energy dispersive X-ray spectroscopy (TEM-EDX)**

The distributions of elements within the samples were qualitatively determined according to the method conducted by Sheraz et al. [52]. In detail, the mycelia or trapping devices were placed in a primary fixative consisting of 2.5% (v/v) glutaraldehyde for approximately 6 hours at 4 °C. The samples were then washed three times at 15-minute intervals with phosphate-buffered saline (PBS) buffer (0.1 mol/L, pH 6.8), followed by post-fixation in 1% (m/v) osmium tetroxide for 2 hours at 4 °C. After washing with PBS buffer, the samples were dehydrated through a graded ethanol series: 30% (v/v) for 15 minutes, 50% (v/v) for 15 minutes, 70% (v/v) for 15 minutes, 80% (v/v) for 15 minutes, 90% (v/v) for 15 minutes, and 100% (v/v) for 15 minutes, twice. Prior to embedding, the samples were infiltrated with a mixture of anhydrous ethanol and epoxy resin EPON-812 (1:1, v/v) overnight. The samples were then placed in an oven at 60 °C for 48 hours and finally stored in desiccators.

Structures were further sectioned into ultrathin slices using an ultramicrotome (Leica RM2016, Germany). The resulting 100-nm ultrathin sections were mounted on 200-mesh copper TEM grids (Agar Scientific). Where indicated, sections were stained with 1% uranyl acetate for 15 minutes, followed by Reynold’s lead citrate (Reynolds, 1963) for 7 minutes. Samples were then observed using a JEOL 1400 transmission electron microscope (JEOL, Akishima, Japan) at 80 kV. To qualitatively determine the distribution of elements within the samples, stained sections were analyzed to identify Fe deposits and quantify Fe content through X-ray microanalysis. Energy dispersive X-ray spectroscopy (EDAX, USA), equipped with the TEM, was employed for this purpose. Circular regions of interest were selected for energy-dispersive X-ray (EDX) analysis. An electron beam with an acceleration voltage of 200 kV and a spot size of 10 nm was focused on the specimen, and X-ray spectra were collected for approximately 100 seconds. X-rays were detected using an Oxford INCAx-sight detector (Oxford Instruments, Abingdon, UK) and analyzed with the Oxford INCA SUITE v.4.02. The Zeta factor method was used for standardless quantitative analysis, checking elements manually selected and reporting elements excluded from qualitative analysis.

**Genome data collection and quality assessment**

To investigate the distribution of a genomic element within the fungal kingdom, we downloaded all published genome and corresponding annotation files from the JGI Mycocosm database (15 December 2022) [53]. Additionally, we used “Orbiliomycetes” as a search term in NCBI’s Genome Browser to retrieve all genome assemblies with gene annotation files for this class. In total, 2,175 fungal genomes were collected. For each predicted gene, the longest transcript was extracted as the representative transcript. Protein sequences for each genome were generated using gffread (v.0.9.12) [54]. To assess the quality of the gene sets for use in phylogenomic analyses, we evaluated their completeness using Benchmarking Universal Single-Copy Orthologs (BUSCO) (v.3.0.2) [55] based on the fungi_odb10 lineage dataset (2021-06-28). Genes were categorized as “single copy” if only one complete predicted gene was present in the genome, “duplicated” if two or more complete predicted genes were found for one BUSCO gene, “fragmented” if the predicted gene was shorter than 95% of the aligned sequence lengths from 50 different fungal species, and “missing” if no predicted gene was present. Only genomes with a BUSCO completeness of 90% or higher were considered high-quality and retained. In total, 2,057 high-quality fungal genomes were retained (Table S3). A maximum likelihood (ML) tree based on concatenated amino acid sequences from 750 single-copy orthologs was constructed.

**Identification homologue of the gene cluster of desferriferrichromes biosynthesis in the** **fungal kingdom**

The biosynthetic gene clusters for each genome were annotated using antiSMASH 7.0 (v.7.0.0) [56] via the command-line version with default parameters, except for the argument ‘--taxon fungi’. In total, 67,350 secondary metabolism gene clusters were annotated from 1,488 fungal genomes. A local database was created from GenBank sequences of all gene clusters using the ‘cblaster makedb’ module (v.1.3.18) [57]. To identify gene clusters homologous to the desferriferrichrome gene cluster within the fungal kingdom, the search function of cblaster was employed to query the amino acid sequences of the seven genes in the desferriferrichrome gene cluster (*Ao410–Ao416*) against the local database. Both *Ao414* (*sidA*) and *Ao415* (*NRPS*) needed to be represented in a hit cluster, and the minimum percent identity for a BLAST hit was set to 30%. There are 927 fungal genomes containing desferriferrichrome gene cluster homologues, all of which were from Ascomycetes. The presence and absence of the desferriferrichrome gene cluster were mapped onto a phylogenetic tree (containing 2,075 high-quality fungal genomes) to visualize the distribution of the desferriferrichrome gene cluster within the fungal kingdom.

**Inference of homologous groups**

To better understand the evolutionary trajectory of the desferriferrichromes biosynthetic gene cluster within the fungal kingdom, we selected 22 representative fungi (9 Orbiliomycetes, 4 Pezizomycetes, 3 Sordariomycetes, 2 Eurotiomycetes, 2 Saccharomycetes, and 1 Taphrinomycetes) containing the desferriferrichromes gene cluster, as well as 1 fungus from Cystobasidiomycetes lacking the desferriferrichromes gene cluster, to serve as an outgroup. This selection was guided by the fungal species tree from the MycoCosm portal. Protein sequences from these 22 fungal genomes, totaling 131,694 sequences, were subjected to all-versus-all BLASTP (v.2.11.0+) [58]. The resulting output was then analyzed using OrthoFinder (v.2.5.2) [59] for gene family clustering, employing the MCL algorithm with an inflation factor set to 1.5. This analysis identified 1,136 sets of 1:1 orthologous protein-coding genes.

**Phylogenetic analyses and divergence times estimating**

Amino acid sequences for each orthogroup were aligned using MAFFT (v.7.505) [60] with the ‘auto’ parameter. Codon alignments were generated from these MAFFT-aligned amino acid sequences using PAL2NAL (v14) [61] with default parameters. The aligned amino acid sequences and their corresponding codon alignments were concatenated to construct amino acid (AA) and nucleotide (NT) data matrices, respectively. For the AA data matrix, the best substitution model was determined using the “-m TEST” parameter, which estimates the optimal model based on Bayesian Information Criterion (BIC) values. Maximum likelihood (ML) trees with 1,000 bootstrap iterations were constructed according to the best-fitting model using IQ-TREE2 [62]. Fourfold degenerate sites were extracted from the NT data matrix using the open-source tool get4foldSites (https://github.com/brunonevado/get4foldSites). Divergence times for all nodes were estimated with MCMCTree from the PAML package [63], using 275,133 fourfold degenerate sites from 1,136 single-copy orthologous genes. Calibration was based on three major events [30]: Ascomycota crown group (487–773 Mya), Orbiliomycetes–other Pezizomycotina (353–554 Mya), Saccharomycotina crown group (276–514 Mya) and one fossil calibration point (A extinct NTF group with ring trapping structure: >100 Mya) [31]. The clock model was set to independent rates, with a burn-in of 20,000, 100,000 samples, a sampling frequency of 10, and the JC69 model. Tracer v.1.7.1 [64] was used to visually check the convergence of parameters across replicate MCMC chains. All resulting effective sample sizes (ESS) ranged from 275 to 1,299 (Table S6), indicating that the parameters across replicate MCMC chains achieved convergence (with ESS values expected to exceed 200).

**RNA-seq data analysis**

Raw reads from both WT and Δ*SidA* samples were examined using FastQC. To obtain clean data, fastp (v.4.10.0) [65] was used with default parameters to filter adaptor sequences and remove low-quality reads. The reference genome and gene model annotation files for A. oligospora ATCC 24927 were downloaded from NCBI’s Genome Browser. Genome indexing was performed using ‘hisat2-build’ in Hisat2 (v.2.2.1) [66], incorporating splice sites and exon information. The filtered paired-end reads were aligned to the reference genome using Hisat2. Gene abundance for each sample was estimated using featureCounts (v.2.0.3) [67] from the aligned BAM files. Differential expression analysis was conducted using the DESeq2 R package (v.1.30.1) [67]. DESeq2 employs statistical tests to identify differential expression in digital gene expression data based on the negative binomial distribution. Resulting p-values were adjusted using the Benjamini–Hochberg approach to control the false discovery rate.

**Temperature and metal ion assays**

Conidial suspensions (50 μL) were collected and quantified using a hemocytometer. Subsequently, 100 μL of fungal spore suspensions (800 spores/μL) were added to 6 cm Petri dishes containing water agar (20 g/L agar in 1 L double-distilled water (ddH_2_O). Water agar prepared with tap water (Kunming Water Supply Company) was used as a negative control. All fungal dishes were incubated at 12 °C, 20 °C, and 28 °C for 7 days. Trapping devices on each plate were observed and counted under a microscope. For colony growth analysis, fungal strains were cultivated on 9 cm diameter PDA plates at 12°C,16 °C, 20 °C, 24 °C, 28°C and 32 °C for 6 days, and the diameter of each colony was measured. All experiments were conducted with at least three replicates.

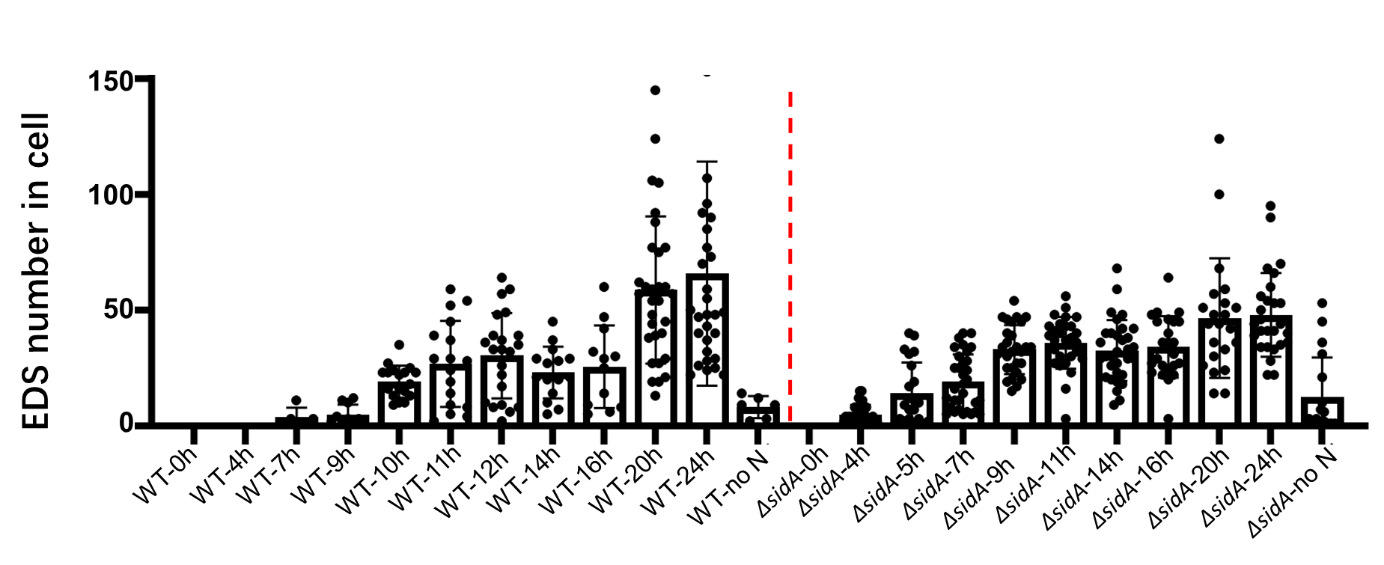

**Fig. S1 Comparison of the formation time of electron-dense bodies in WT and *ΔsidA* mutant strains.**

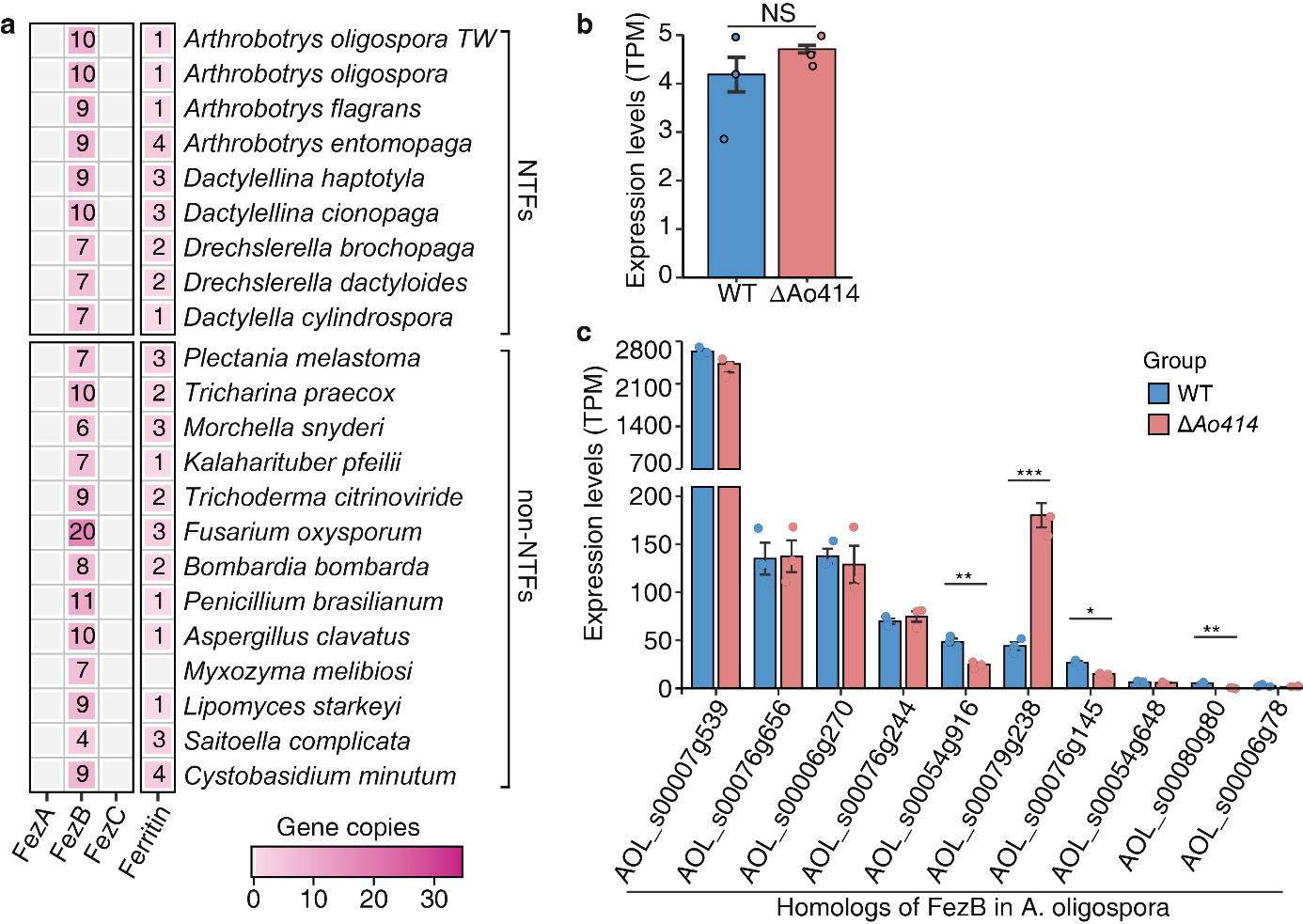

**Fig. S2 Homologs of iron storage related genes in nematode-trapping fungi.** **(a)**, Distributions of ferrosomes related genes and ferritin in NTFs and non-NTFs. **(b)**, Expression levels of ferritin homologs in WT and *Ao414* mutant of *A. oligospora.* **(c)**, Expression levels of FezB homologs in WT and *Ao414* mutant of *A. oligospora.*

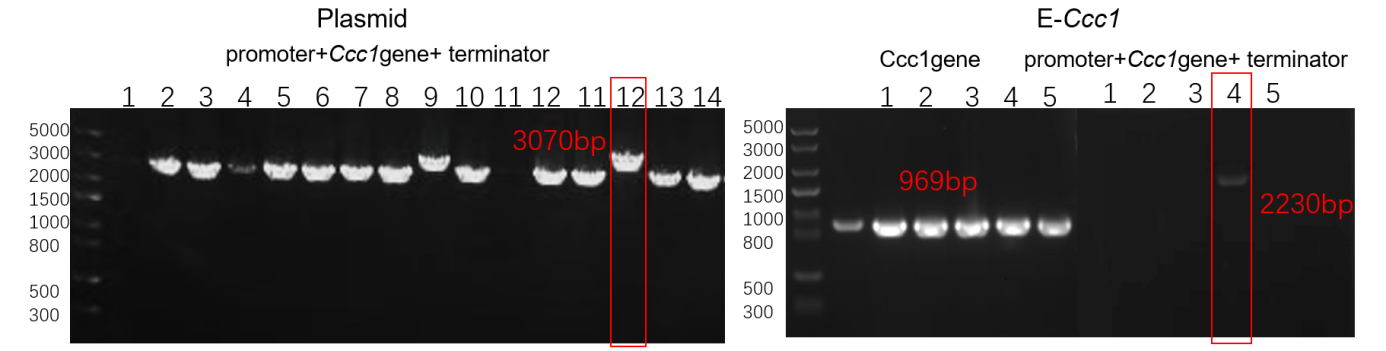

**Fig. S3 Construction and PCR of mutants E-*Ccc1*.**

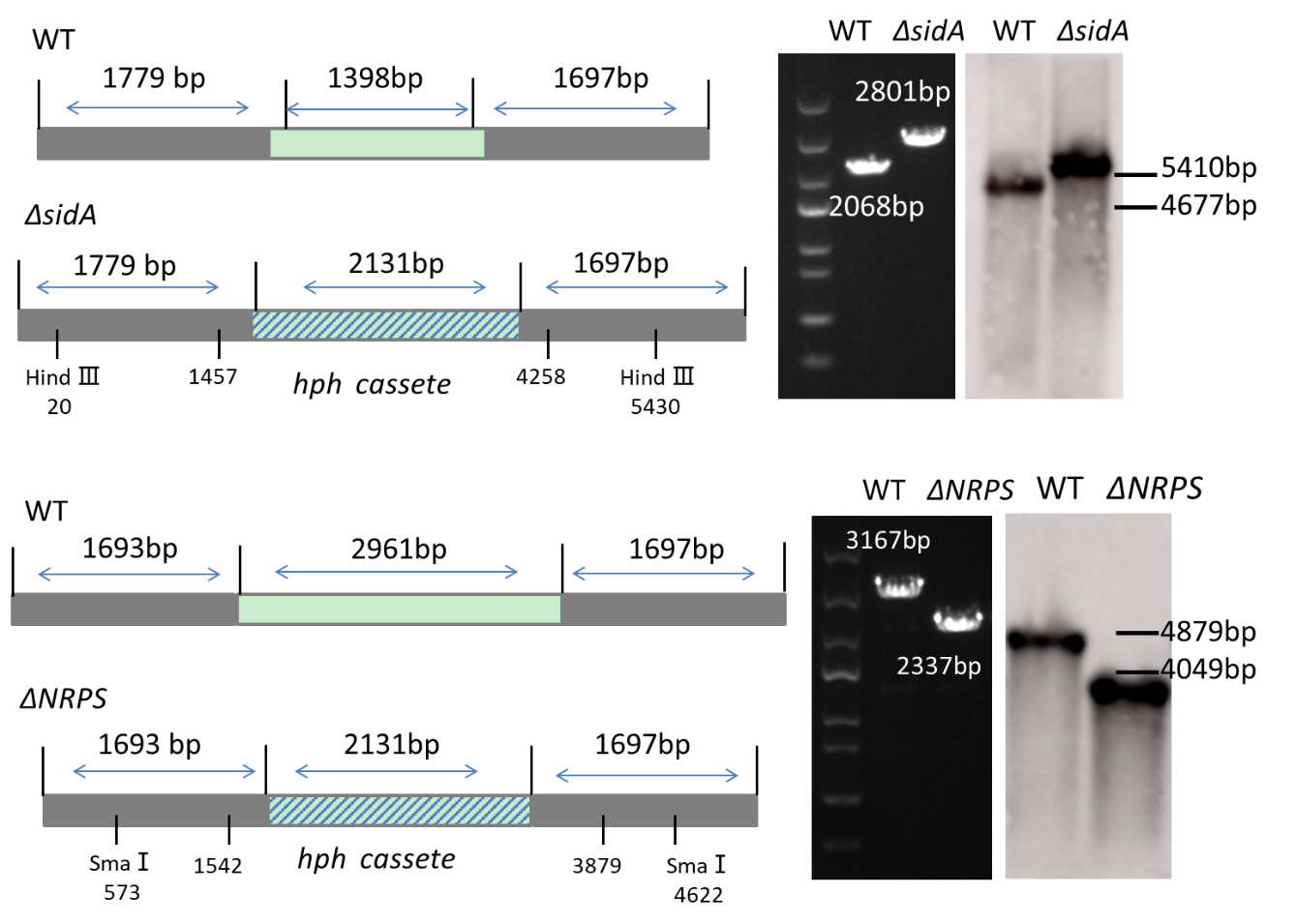

**Fig**. **S4** **Construction and PCR and Southern blot analysis of two mutants Δ*sidA* and Δ*NRPS*.**

**Table S1. Evaluation of iron distribution and contents in trapping devices of *A. oligospora*.** Energy Dispersive X-ray Spectroscopy (EDX) analysis of in the trapping devices of WT and *ΔsidA*.
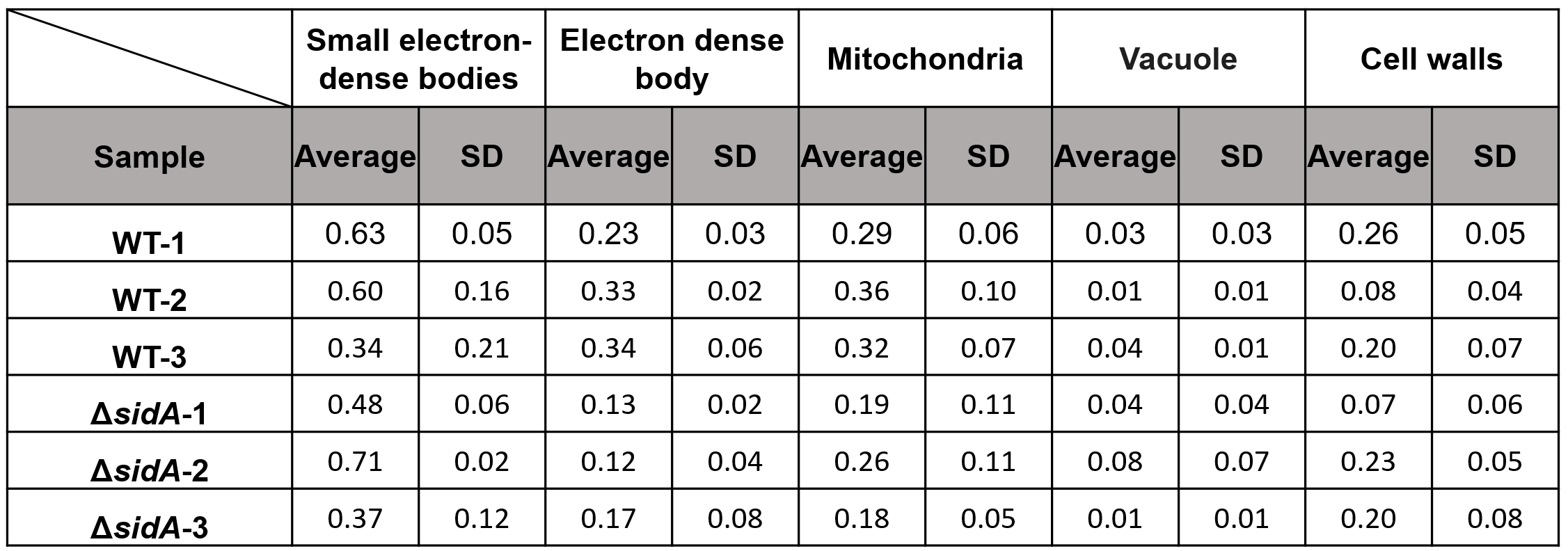

**Table S2. Evaluation of the functions of desferriferrichrome in iron distribution and storage in *A. oligospora* mycelia.**  Energy Dispersive X-ray Spectroscopy (EDX) analysis of WT mycelia treated with only solvent DMSO, WT mycelia treated with desferriferrichrome, *ΔsidA* mycelia treated with only solvent DMSO and *ΔsidA* mycelia treated with desferriferrichrome.

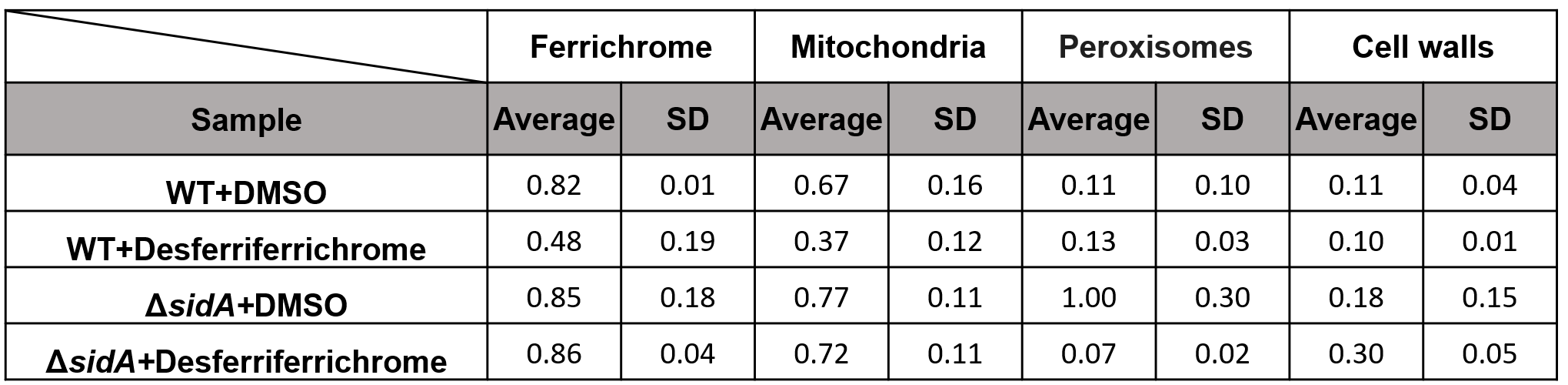

**TableS4.** **The primers were used for construction of the mutants Δ*sidA* and Δ*NRPS*.**

| **Gene** | **Primer name** | **Sequence (5' - 3')** |
| --- | --- | --- |
| *sidA* | *sidA-*UP-F | CGAGGGGACTGCCTTGAAG |
|  | *sidA-*UP-R | CTCCTTCAATATCATCTTCTGTCGAGGGACTCAACGCTGTATTGAC |
|  | *sidA-*Down-F | CACTTGTTTAGAGGTAATCCTTCTTCCTTGCCGTGCGTGCAGG |
|  | *sidA-*Down-R | TAGAGGTGAACGGTCACCCAGTC |
|  | *sidA-*YZ-F | CGGCAAGATCAAGGAGGC |
|  | *sidA-*YZ-R | GGATCGAGCCAGTAGAAATCCA |
| *NRPS* | *NRPS-*UP-F | GAATAGCAGCATATCGACGGA |
|  | *NRPS-*UP-R | CTCCTTCAATATCATCTTCTGTCGAAGACTCTCCGTCAATCTGTAT |
|  | *NRPS-*Down-F | CACTTGTTTAGAGGTAATCCTTCTTCTCTTGGAAGAAGGGCGG |
|  | *NRPS-*Down-R | CATCACCCAAATCCATGGC |
|  | *NRPS-*YZ-F | GGTTCAGAGACAGAGCTCTTG |
|  | *NRPS-*YZ-R | CCGATGGTTTCTAGCTCTGAAC |
| Hyg | Hyg-F | TCGACAGAAGATGATATTGAAGGA |
|  | Hyg-R | AAGAAGGATTACCTCTAAACAAGT |

**Table S5.** **The primers were used for construction of mutant E-*Ccc1*.**

| **Gene** | **Primer name** | **Sequence (5' - 3')** |
| --- | --- | --- |
| Ccc1 | *Ccc1*-cDNA-F | cctattctacccaagcatcgatATGTCCATTGTAGCACTAAAGAAC |
|  | *Ccc1*-cDNA-R  *Ptrpc*-F  *Ptrpc*-R  *Ccc1*-yz-F  *Ccc1*-yz-R  *Ccc1P*-RT-F  *Ccc1P*-RT-R | CACGAGCTACTACAGATCCCCGTTAACCCAGTAACTTAACAAAGAACC  CGGGGATCTGTAGTAGCTCGTG  atcgatgcttgggtagaatagg  gcacaggtacacttgtttag  CAGCGTGAGCTATGAGAAAG  TCTAGACGAACCTGCCGAGA  ACACCACCCACAACAACCAT |

**Table S6. Effective sample sizes (ESS) of the convergence of parameters across replicate MCMC chains.**

| Statistic | Mean | ESS | Type |
| --- | --- | --- | --- |
| t_n27 | 4.129 | 275 | R |
| t_n30 | 2.202 | 277 | R |
| t_n28 | 2.578 | 280 | R |
| t_n35 | 1.124 | 296 | R |
| t_n37 | 3.377 | 323 | R |
| t_n38 | 3.247 | 326 | R |
| t_n34 | 1.558 | 337 | R |
| t_n40 | 1.279 | 339 | R |
| t_n43 | 3.616 | 348 | R |
| t_n36 | 3.703 | 356 | R |
| t_n31 | 0.859 | 356 | R |
| t_n29 | 2.429 | 421 | R |
| t_n32 | 0.157 | 435 | R |
| t_n26 | 4.635 | 447 | R |
| t_n23 | 6.445 | 466 | R |
| t_n33 | 1.934 | 534 | R |
| t_n39 | 2.758 | 584 | R |
| t_n24 | 5.165 | 750 | R |
| t_n25 | 4.885 | 771 | R |
| t_n42 | 1.77 | 1265 | R |
| t_n41 | 2.213 | 1299 | R |
| mu | 0.336 | 552 | R |
| sigma2 | 0.027 | 979 | R |
| kappa | 2.620 | 16879 | R |
| lnL | -2622000 | 17304 | R |
